# Calibration of Agent Based Models for Monophasic and Biphasic Tumour Growth using Approximate Bayesian Computation

**DOI:** 10.1101/2022.09.13.507714

**Authors:** Xiaoyu Wang, Adrianne L. Jenner, Robert Salomone, David J. Warne, Christopher Drovandi

**Affiliations:** School of Mathematical Sciences, Queensland University of Technology, Brisbane, QLD, Australia; Centre for Data Science, Queensland University of Technology, Brisbane, QLD, Australia; School of Computer Science, Queensland University of Technology, Brisbane, QLD, Australia

**Keywords:** Agent-based model, Cancer, Likelihood-free inference, Parameter estimation, Tumour growth

## Abstract

Agent-based models (ABMs) are readily used to capture the stochasticity in tumour evolution; however, these models are often challenging to validate with experimental measurements due to model complexity. The Voronoi cell-based model (VCBM) is an off-lattice agent-based model that captures individual cell shapes using a Voronoi tessellation and mimics the evolution of cancer cell proliferation and movement. Evidence suggests tumours can exhibit biphasic growth *in vivo*. To account for this phenomena, we extend the VCBM to capture the existence of two distinct growth phases. Prior work primarily focused on point estimation for the parameters without consideration of estimating uncertainty. In this paper, approximate Bayesian computation is employed to calibrate the model to *in vivo* measurements of breast, ovarian and pancreatic cancer. Our approach involves estimating the distribution of parameters that govern cancer cell proliferation and recovering outputs that match the experimental data. Our results show that the VCBM, and its biphasic extension, provides insight into tumour growth and quantifies uncertainty in the switching time between the two phases of the biphasic growth model. We find this approach enables precise estimates for the time taken for a daughter cell to become a mature cell. This allows us to propose future refinements to the model to improve accuracy, whilst also making conclusions about the differences in cancer cell characteristics.

## 1 Introduction

Cancer is a disease that arises through the progressive alteration of normal cells. Solid cancers form what is commonly referred to as tumours, which are populations of cancerous cells joined together with connective tissue (Weinberg and Weinberg, 2006). Over time, these tumours expand and take up residency in healthy tissue. To elucidate the mechanisms of the cancer pathophysiology, researchers have mainly focused on the analysis of aberrant cell dynamics within tumour structures. The pivotal role these abnormal cells play in oncogenesis constitutes a salient focus of these investigations (Noble, 2002; Markowetz, 2017).

Mathematical modelling of cancer growth and development has been used to study the dynamical process of cancer cells for many years (Altrock et al., 2015; Beerenwinkel et al., 2015; Barbolosi et al., 2016; Tabassum et al., 2019). Deterministic approaches, such as systems of ordinary differential equations (ODEs) and systems of partial differential equations (PDEs), have been used successfully to model cancer growth (Villasana and Radunskaya, 2003; Yafia, 2011; Tao et al., 2014; Jenner et al., 2020b; Dehingia et al., 2021; Klowss et al., 2022; VandenHeuvel et al., 2022). While insightful, generally these models do not capture the phenotypical and spatial heterogeneity that arises through stochastic processes of tumour growth, or consider the behaviour at an individual cancer cell level (Irurzun-Arana et al., 2020). In turn, such models often do not capture dynamical processes such as individual cell proliferation (Iyer et al., 2011; Sahoo et al., 2011) and cell movement (Groh and Louis, 2010). For instance, Warne et al. (2022) highlight that continuum models need to align with specific assumptions that can diverge from reality. Specifically, standard deterministic continuous models that are typically used to represent cell invasions often require cell motility rates to dominate proliferation rates. Such conditions may not hold true for slow-growing tumours. In such tumours, pressure-driven mobility can serve as the principal catalyst for tumour growth. Consequently, stochastic discrete models provide a more adaptable framework for modelling various spatiotemporal patterns, such as clustering. As a result, parameter estimates relating to the proliferation rate of tumor cells are often underestimates of the true proliferation rate due to spatial clustering inhibiting growth through contact inhibition. By explicitly modelling the proliferation and motility processes of a spatially heterogeneous tumor using an ABM the accurate parameter estimates and realistic uncertainty quantification can be obtained, even in situations with spatially irregular boundaries with fingering patterns or multiple local clusters.

More recently, agent-based models (ABMs) have become a popular approach for capturing the stochastic nature of tumour growth (Ozik et al., 2018; Metzcar et al., 2019; Macnamara, 2021; Klowss et al., 2022). ABMs model cells as agents, and the interactions of cells are governed by probabilistic rules that depend on physically-meaningful parameters (An, 2012). There are two popular paradigms of ABMs: on-lattice and off-lattice models (Railsback and Grimm, 2019). On-lattice models restrict movement of cells to neighbouring points on a pre-defined lattice. Whereas cell movement in off-lattice models are governed by interaction forces between neighbouring cells and the surrounding environment. Although, one drawback of ABMs is that model simulation can be more computationally expensive than solving a deterministic system of differential equations.

This work focuses on investigating and extending the Voronoi cell-based model (VCBM), an off-lattice ABM of tumour growth in 2-dimensions based on the model by Jenner et al. (2020a), with the concept originally introduced by Kansal et al. (2000b,a); Schaller and Meyer-Hermann (2005); Kempf et al. (2010); Fletcher et al. (2013) and more recently investigated by Cleri (2019); Germano et al. (2022). Agents are defined as either healthy cells or tumour cells. The movement of cells through the domain is modelled off-lattice using a force-balance equation based on Hooke’s law to account for cell adhesive and repulsive forces. Cell boundaries are then defined by a Voronoi tessellation.

Many growth factors are known to influence cell behaviour in a biphasic manner, causing an increase or decrease in cellular division contingent upon their concentration (Konstorum et al., 2013). A biphasic relationship has been observed between the speed of an invading tumour front and the concentration of collagen in the surrounding gel, whereby two growth phases can be observed in *in vitro* or *in vivo* tumour growth (Konstorum et al., 2013). This biphasic relationship could be due to the complex interaction between cell proliferation and migration (Perumpanani and Byrne, 1999). This insight led to the development of mathematical models that could test these hypotheses (Marchant et al., 2006; Konstorum et al., 2013).

Marchant et al. (2006) successfully devised a theoretical model of malignant invasion that accurately reproduces the biphasic dependency of tumour cell invasion speed on the density of the surrounding normal tissue. To encapsulate this biphasic phenomenon effectively within the context of the VCBM, we introduce the concept of a switching time, such as in Murphy et al. (2022). This switching time refers to a specific point in time in which the tumour evolution transitions between two distinct growth phases, that is, it reflects transition of tumour growth *in vivo* from an ‘establishment’ phase, characterized by initial proliferation and localization, to an ‘expansion’ phase, where rapid and invasive growth occurs (Marchant et al., 2006). This transition point, derived from *in vivo* measurements, provides a biologically significant depiction of the tumour’s temporal evolution. Thus, such an approach enhances the VCBM’s ability to simulate the complex dynamics of tumour growth accurately.

Cancer cell proliferation is largely the driving factor of tumour growth and is impacted by spatial limitations and nutrient sources (Iyer et al., 2011; Sahoo et al., 2011; Jenner et al., 2020a). As such, the key mechanism of tumour growth in the VCBM is cancer cell proliferation (Cheng et al., 2012; Benzekry et al., 2014). Calibrating the cellular proliferation parameters to tumour growth measurements would provide a better understanding of the differences between tumour types. However, obtaining parameter estimates for an ABM is not trivial. This work proposes a Bayesian framework to calibrate cancer cell proliferation in the VCBM and capture the tumour growth in ovarian, pancreatic and breast tumour cell lines implanted *in vivo*.

ABMs are simulation-based models, for which inference can be challenging as the likelihood function for such models is often intractable. Hence, approximate Bayesian computation (ABC) (Sisson et al., 2018) is proposed to bypass the likelihood function and estimate the model parameters. ABC requires that the model can be simulated, but does not require evaluation of its corresponding likelihood function. ABC has been previously used to calibrate a small number of biological ABMs. For example, Lambert et al. (2018) apply ABC to a lattice model and Ross et al. (2017) use ABC to improve the experimental design of a wound-healing assay by calibrating a cell-cell adhesion model (Khain et al., 2007; Ross et al., 2015). Rocha et al. (2021) use ABC to calibrate to identify cell motility parameters using biological experiments in their ABM developed in PhysiCell (Ghaffarizadeh et al., 2018). However, they use simple ABC algorithms which require a large number of model simulations, and are thus computationally expensive when the model simulation is not computationally trivial. We calibrate the VCBM to tumour growth time series data using sequential Monte Carlo (SMC)-ABC, specifically the algorithm of Drovandi and Pettitt (2011), since it is much more efficient than the standard ABC rejection algorithm, and it can easily take advantage of parallel computing resources.

Understanding the limitations and flexibility of the VCBM is important to be able to determine its reliability as a predictor of tumour growth. The main contribution of this paper is to provide an efficient Bayesian approach for fitting off-lattice stochastic models of tumour growth to time series data of tumour volumes. As a case study, we extend the VCBM proposed by Jenner et al. (2020a) to a biphasic VCBM, which has not been previously calibrated to data in the literature. By using an efficient Bayesian algorithm, we are also able to thoroughly investigate if the VCBM is flexible enough to recover real tumour volume time series data across a range of cancers. Specifically, we calibrate the model to *in vivo* measurements for breast cancer, ovarian cancer and pancreatic cancer in mice obtained from Wade (2019); Wade et al. (2020) and Kim et al. (2011). Calibration results suggest that subcutaneous growth leads to less spatial inhibition as the tumour grows. Biomechanically, this suggests that the growth dynamics evolve alongside the tumour’s progression. Subcutaneously grown tumours are known for exhibiting negligible propensity for metastasis due to a lack of spatial constraints, which is one of the causes of metastases associated within an intra-organ growth (Schmidt et al., 2016). Furthermore, it is apparent that cellular proliferation changes over time due to potential cellular mutations that result in faster or slower cell proliferation (Bozic et al., 2010) or even necrosis from vascular limitations. Our calibration approach reveals that the model is sufficiently robust in capturing both monophasic and biphasic growth occurring across data sets.

We now describe the structure of the paper. In Section 2, we describe the data we analyse, explain the VCBM that we calibrate to the data and provide details of the ABC calibration method that we adopt for fitting the VCBM to data. Section 3 presents the results of the calibration process. In Section 4, we discuss the findings further, the limitations of our study, and directions for future research.

## 2 Methods

### 2.1 Experimental measurements: *in vivo* tumour volume

Measurements from three independent *in vivo* cancer datasets (Fig 1) are used to calibrate parameters in the VCBM: breast cancer cells (Kim et al., 2011), ovarian cancer cells (Kim et al., 2011), and pancreatic cancer cells (Wade, 2019; Wade et al., 2020). In these experiments, tumour volume (*mm*^3^) is approximated using caliper measurements in two-dimensions:

**Fig. 1:**
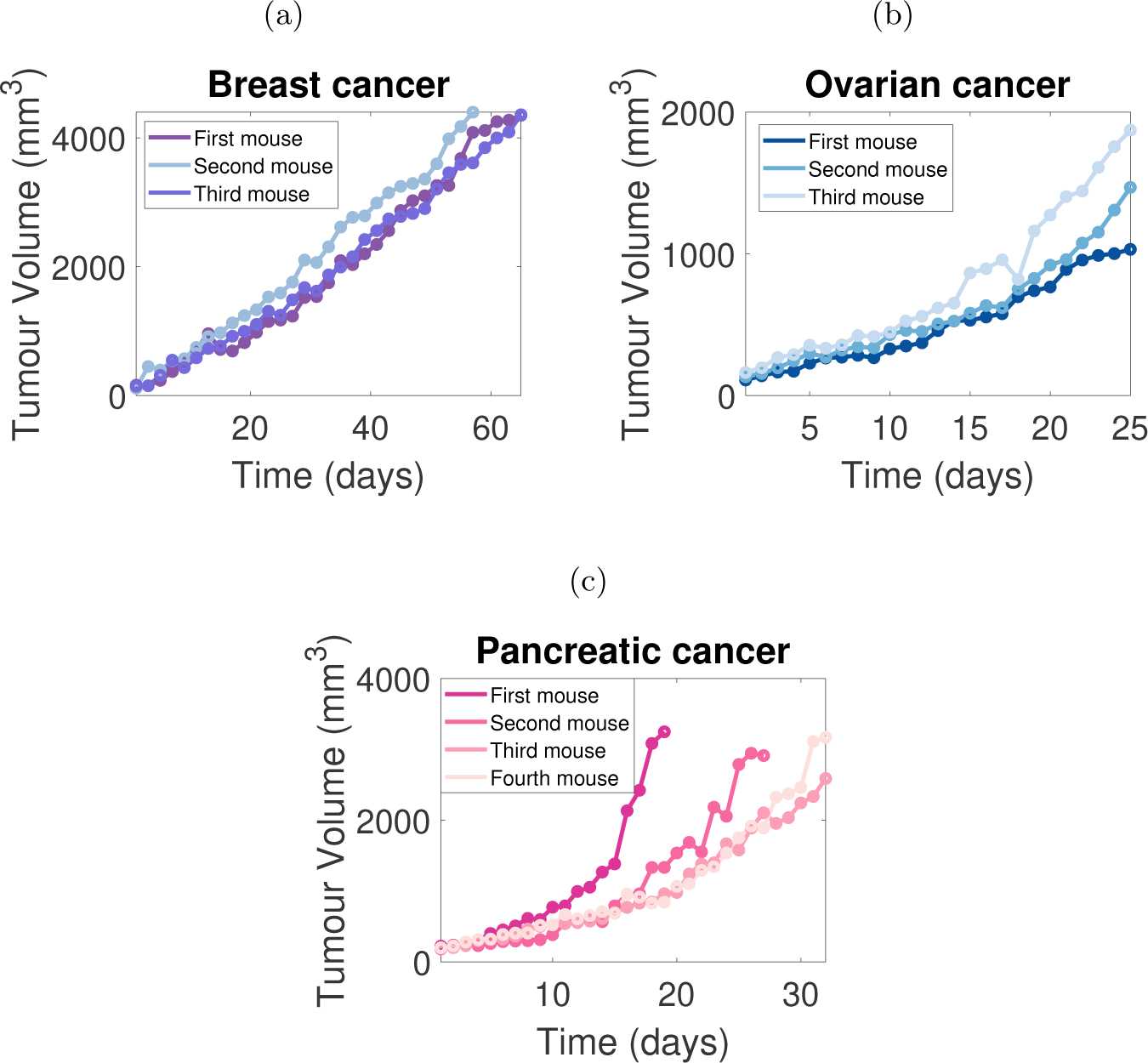
*In vivo* tumour volume measurements. Tumour growth is measured *in vivo* for (a) breast cancer cells (*n* = 3) (Kim et al., 2011), (b) ovarian cancer cells (*n* = 3) (Kim et al., 2011) and (c) pancreatic cancer cells (*n* = 4) (Wade, 2019; Wade et al., 2020). Each mouse is identified by a unique colour and the specific data measurements are represented by solid points. The ordering of the naming convention for the mice was chosen randomly. Pancreatic tumour datasets in (c) exhibit a potentially biphasic tumour growth pattern.

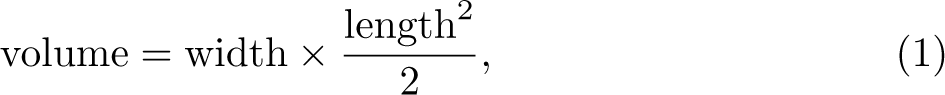

where width (*mm*) is the longest tumour measurement and length (*mm*) is the tumour measurement along a perpendicular axis (Kim et al., 2011). We also generate a synthetic dataset designed to mirror the quantity of data points present in experimental data. This approach enables an evaluation of the calibration method’s ability to accurately recover known parameter values. The synthetic and experimental datasets are briefly described below. Full details of the experimental data can be found in Kim et al. (2011) and Wade (2019); Wade et al. (2020).

#### 2.1.1 Breast cancer data

Kim et al. (2011) measure the *in vivo* breast cancer tumour volume over 66 days, with measurements being recorded every second day. Each mouse is injected with Her2/neu-expressing human breast cancer cells MDA-MB435, and tumour volume measurements commenced when the seeded tumours reach 100 − 120 *mm*^3^. Fig 1a depicts tumour growth data for four mice with breast cancer. We remove measurements from mouse two from day 58 onward as the tumour volume began to exhibit a decline, which we deem biologically infeasible for our chosen model assumptions. For more details, see Kim et al. (2011).

#### 2.1.2 Ovarian cancer data

Kim et al. (2011) measure the *in vivo* ovarian cancer tumour volume over 25 days, with measurements being recorded daily in mice injected with SK-OV3 cells. The tumour volume measurements commence when the seeded tumours reach 100 − 120 *mm*^3^. Fig 1b displays tumour growth data for three mice with ovarian cancer. For more details, see Kim et al. (2011).

#### 2.1.3 Pancreatic cancer data

Tumour growth measurements in mice seeded with pancreatic cancer cells Mia-PaCa-2 are collected by Wade (2019); Wade et al. (2020). Fig 1c displays the four lines representing the tumour volume for four different mice over 33 days which were recorded every day. The cessation of tumour volume measurements occurred once the volume exceeded a given threshold. Hence, the final observation time varies between mice.

#### 2.1.4 Synthetic cancer data

In this study, we generate three synthetic datasets using the model with predetermined parameter values, as outlined in Table S1. Specifically, we generate three synthetic cancer datasets utilizing a biphasic model with a fixed switching time. To ensure consistency with actual datasets that exhibit biphasic growth, which we observe to have a measurement length of 32 days, we set the length of these three synthetic datasets to 32 days. Additionally, we generate two synthetic datasets demonstrating monotonic exponential growth in order to validate the original VCBM.

### 2.2 Voronoi Cell-based Model (VCBM)

ABMs are popular computational models that model cells as agents with associated probabilistic sets of rules for the purpose of capturing stochastic cellular dynamics. In this study, we model tumour growth using an ABM as it allows us to capture the spatial evolution of a tumour. In addition, ABMs enable simulation of an array of stochastic processes and can account for individual cell behaviours, including low movement rates, and their interactions within the surrounding microenvironment. This is a more detailed description of the tumour compared to an analogous deterministic system (e.g. ODE or PDE). To enable flexibility in cellular movement processes, we use an off-lattice ABM. Although on-lattice ABMs (Lundh, 2007; Lowengrub et al., 2009; Cristini and Lowengrub, 2010; Voss-Böhme, 2012; Wang et al., 2015; Poleszczuk et al., 2016; Norton et al., 2019) are less expensive to simulate, off-lattice ABMs can be more effective at simulating the heterogeneous nature of the tumour microenvironment.

Here, we describe the VCBM, an off-lattice model developed by Jenner et al. (2020a, 2022) that simulates the 2-dimensional cellular dynamics of tumour formation. In this study, we simplify their model to capture control *in vivo* tumour growth only, omitting the treatment-related agents of virusinfected cancer cells, dead cancer cells, and empty space. As a result, we only investigate three primary dynamics governing tumour growth: cancer cell proliferation, cell movement, and cell invasion into healthy tissue. We assume that no cells die during the simulation and that healthy cells are pushed outward solely by the force of multiplying cancer cells.

#### 2.2.1 Voronoi tessellation

The 2-dimensional spatial position of the *i*th cell agent at time *t* is defined by a center point ***r****_i_*(*t*) ∈ ℝ^2^. We consider a lattice as the set of cell agent points represented by the vector ***r***(*t*), where the size of ***r***(*t*) depends on the number of cells in the domain at time *t*. We consider the area of a cell to be the region of space enclosed by a Voronoi tessellation formed by the cell centre point. A Voronoi tessellation is used to define the edges of a cell (see Fig 2), i.e. the boundary of cell *i* is the line equidistant between the cell’s centre point and one of its connected cell’s centre points. Voronoi cells on the boundary have an infinite area, and these cells are not explicitly modelled as contributing to the VCBM dynamics. Compared to on-lattice models, the VCBM does not require fixed cell positions. We define the *i*th cell’s neighbourhood, *N* (*i*), as the set of cells connected to that cell by a Delaunay trangulation (Van Liedekerke et al., 2015).

**Fig. 2:**
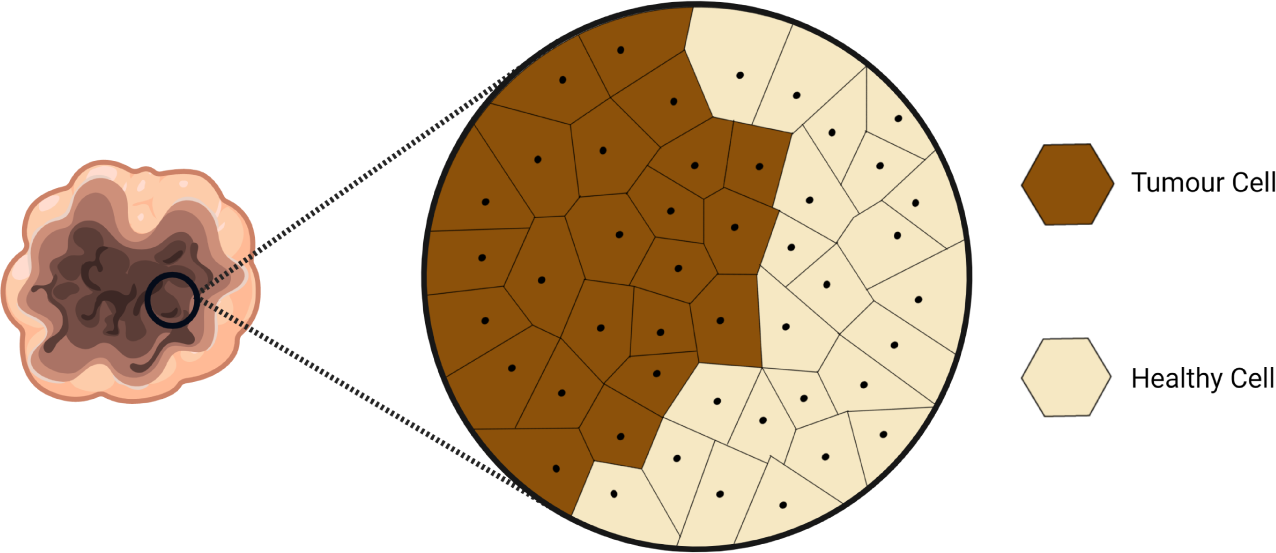
An illustration of the Voronoi tessellation used to model the cell’s shape and its neighbourhood: the boundary of cell *i* is the line equidistant between the cell’s center point (black dot) and one of its connected cell’s center points. Cells in the neighbourhood of a particular cell are those sharing a common edge. A Delaunay trangulation is used to determine neighbouring cells connected to cell *i*.

#### 2.2.2 Cancer cell movement

In the VCBM, cancer cell proliferation is the primary driver of cell movement. The network of forces interacting on cell *i* is modelled by Hooke’s Law (Fig 3a), which computes the effective displacement of the cell and we use this to update the cell’s position. The force between cell *i* and its connected neighbour cell *j* is modelled by a damped spring (Fig 3b), and the spring connecting cell *i* and cell *j* has a rest length s*_i,j_*(*t*), where we assume the rest length can be a function of time *t*. The displacement of the *i*th cell is given by

**Fig. 3:**
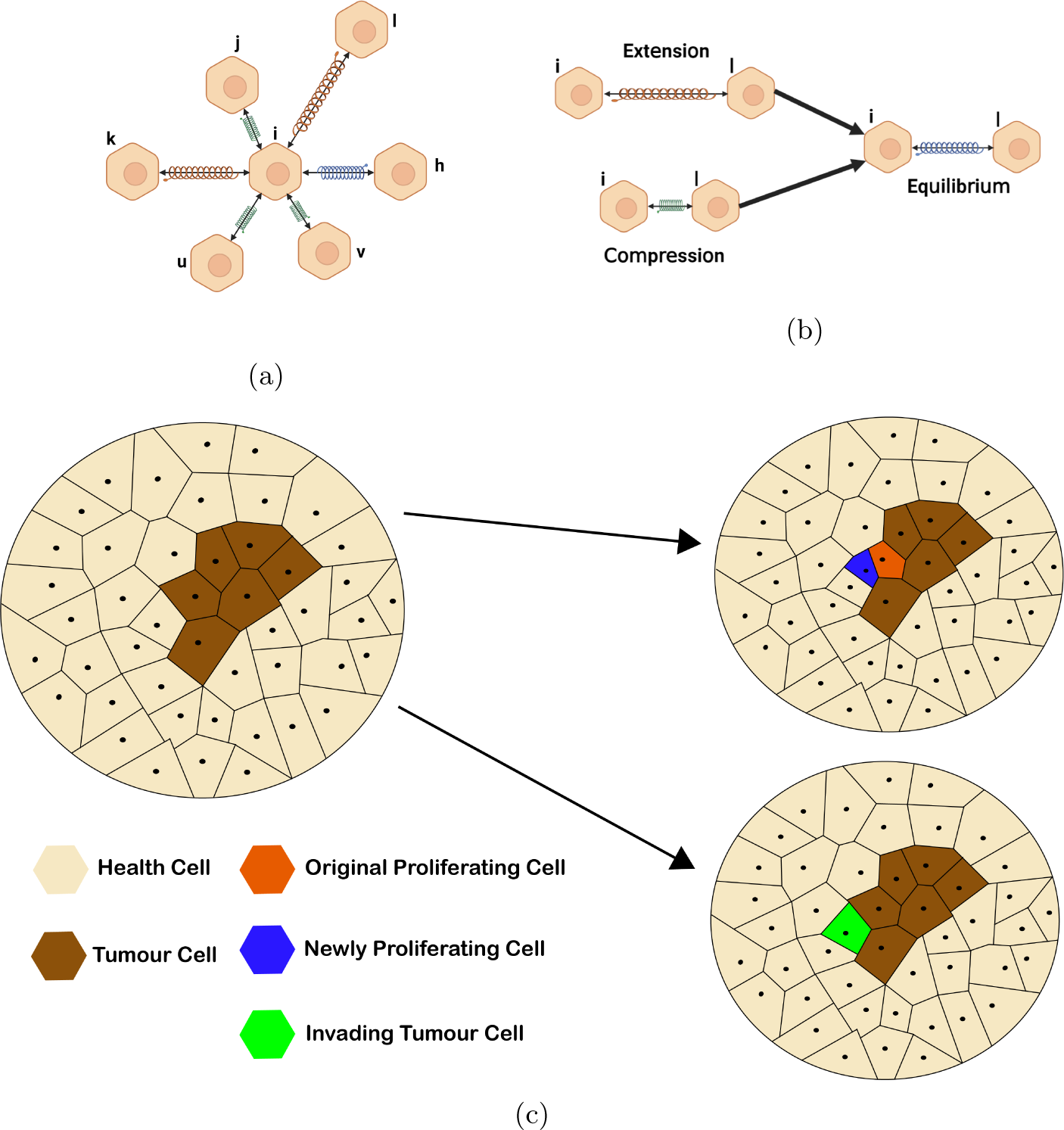
Cancer cell proliferation: (a) the network of forces interacting at a cell *i* is governed by Hooke’s Law, see Eq 4; (b) Hooke’s law models the motion of connected springs as they are extended and compressed and captures how these springs return to a rest state or equilibrium. In the VCBM, we model cell movement using such forces; (c) (upper) the orange cell represents the newly proliferated cell with the blue cell being the original proliferating cell, (lower) the green cell represents to an invading tumour cell that has taken the place of a healthy cell.

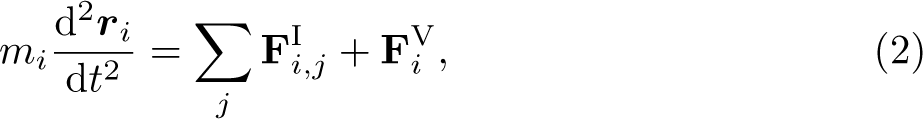

where *m_i_*is the mass of cell *i*, ***r****_i_* is the position of *i*th cell’s centre point, 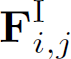 is the interaction force between a neighbourhood cell *j* connected to cell *i* and 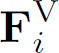 is the viscous force acting on cell *i*.

Following Jenner et al. (2020a); Kansal et al. (2000a,b); Schaller and Meyer-Hermann (2005); Kempf et al. (2010); Fletcher et al. (2013), the total interaction force 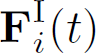 acting on cell *i* at time *t* is given by the sum of the individual forces from the neighbourhood connected cells:

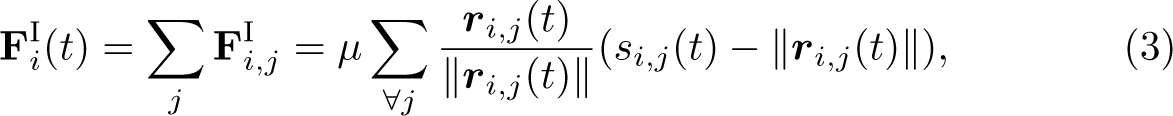

where *µ* is the spring constant, ***r****_i,j_*(*t*) is the vector from the *i*th point to *j*th point at time *t, s_i,j_*(*t*) is the spring rest length between cell *i* and *j* at time *t* and ‖***r****_i,j_(t)*‖ is the Euclidean norm of ***r***_*i,j(t)*_. The viscous force acting on cell *i* is

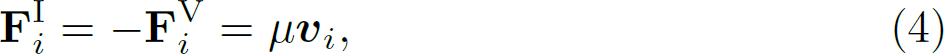

(Galle et al., 2005; Macklin et al., 2012; Jenner et al., 2020a) where ***v****_i_* is the velocity of the *i*th point. By taking a small time interval Δ*t*, the effective displacement of the *i*th point is

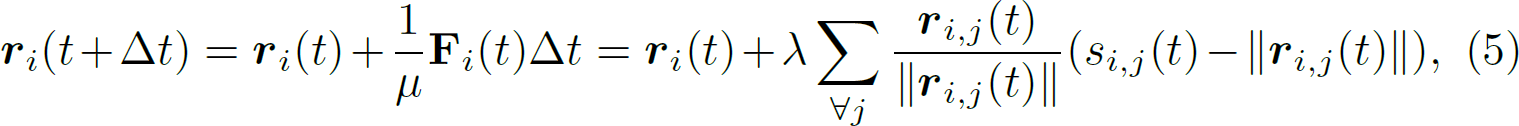

where *µ* is the damping constant. This formulation is readily used in other ABM settings, for example see Murray et al. (2012); Murphy et al. (2019); Browning et al. (2019).

#### 2.2.3 Cancer cell proliferation

For modelling cancer cell proliferation, the Euclidean distance, *d*, between cell *i*’s centre point ***r****_i_*(*t*) and the tumour boundary is used as a proxy for the nutrient source. The boundary of the tumour is defined as the set of tumour cells that reside on the tumour periphery (proliferating edge). We assume cell *i* can proliferate when *d < d*_max_, where *d*_max_ is the maximum radial distance a cell can be from the boundary and still proliferate. The probability of a cell dividing in a given time step Δ*t* is

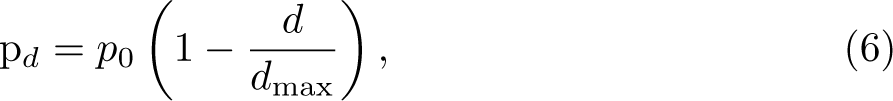

where *p*_0_ is a proliferation constant. In this way, we are defining the probability of proliferation as a function of a cell’s distance to the tumour boundary/nutrient source, as has been done similarly in Kansal et al. (2000a); Jiao and Torquato (2011).

Fig 3c shows that when cell *i* divides it creates two daughter cells *i* and *l*, which are placed at a random orientation equidistant from the original cell *i*’s position. To simulate the growth of a daughter cell to a full grown cell, the resting spring length between cell *i* and cell *l* is initialised as a value less than *s* and increases over time until *s_i,l_* = *s*. Consider *t_d_* is the time since the last cell division, then

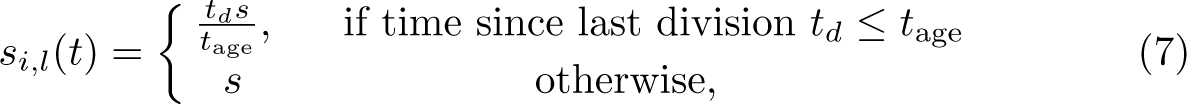

where *s* is the mature resting spring length and *t*_age_ is the time taken for a cell to grow to its full size. In other words, initially at the first time step immediately after division *t_d_* = 1 hour and *s_i,l_*= *s/t*_age_. In the following timestep, the spring length will increase to *s_i,l_*= 2 × *s/t*_age_. The value for *s_i,l_*increases for *t*_age_ time steps, i.e. until *t_d_* = *t*_age_. In this way, we approximate cell growth from a daughter cell size to an adult size which takes *t*_age_ time steps. This variable spring length is what makes the rest spring length a function of time, i.e. *s_i,l_*(*t*). Once it has been *t*_age_ time steps since division we have *s_i,l_*(*t*) = *s*. Another factor for cancer cell proliferation is the time taken for a cell to be able to divide into two daughter cells. In the VCBM, the daughter cells take *g*_age_ time steps to be able to divide again. Note that *t*_age_ *< g*_age_.

#### 2.2.4 Cell invasion

Due to the invasiveness of cancer, we assume a cancer cell’s daughter cell can replace a healthy cell with probability *p*_psc_. In our model, this property refers to the probability that invasiveness occurs at the boundary of tumour tissue. This assumption is inspired by previous ABM work by Jiao and Torquato (2011).

#### 2.2.5 Simulation

The model uses a time-step of one hour, i.e. Δ*t* = 1, which is based off the knowledge that the normal cell cycle is about 24 hours (Bernard and Herzel, 2006) and ovarian cancer’s cell cycle length is greater than 16 hours (Fisi et al., 2016). The VCBM simulation is started by first initialising a square domain with cells arranged in a hexagonal lattice. The cell closest to the centre of the domain is designated as a cancer cell and the remainder are designated as healthy cells. The VCBM is then simulated until it reaches a tumour volume of 100*mm*^2^, which is based off the experiment measurements (Wade, 2019; Kim et al., 2011). Once the tumour reaches this volume, the lattice for healthy and cancerous cells is stored and this lattice is used to simulate the growth of a tumour over the required number of days for each simulation of the model. The model evolves by first checking whether any cancer cell proliferates using Equation 6. For cancer cells that aren’t proliferating in that timestep, they are checked for whether they differentiate into an invasive cell using *p*_psc_. Following this, all cells (healthy, cancerous) are moved using Hooke’s law, Equation 5. Comprehensive descriptions of the model simulation are available in (Jenner et al., 2020a, 2022).

#### 2.2.6 Biphasic model

We find for some datasets that the rate of tumour growth appears to change over time (as shown in Fig 1c). Here, we use a biphasic model to model the change of growth dynamics. To investigate the biphasic tumour growth phenomenon we assume at time *t* = *τ* days, that the parameters can change value. We assume that there are four parameters, i.e. ***θ*** = (*p*_0_*, p*_psc_*, d*_max_*, g*_age_) in the VCBM. In the biphasic version of the model, for *t < τ*, the VCBM is governed by the parameters 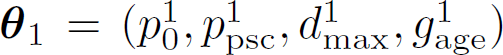), whilst for *t > τ* the VCBM is governed by the parameters 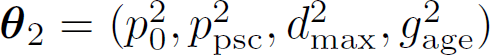). If the tumour growth is monophasic, there are four unknown parameters ***θ***; if it is biphasic, there are nine unknown parameters (***θ***_1_, ***θ***_2_*, τ*).

### 2.3 Bayesian Inference with intractable likelihood

In Bayesian inference, the *posterior distribution* of the parameters ***θ*** is given by

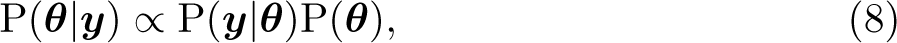

where the *prior distribution* P(***θ***) is updated by the observation data ***y*** = (*y*_1_*, y*_2_*, …, y_T_*) with length *T* via the *likelihood function* P(***y***|***θ***). In this paper, the parameters for the monophasic VCBM are ***θ*** = (*p*_0_*, p*_psc_*, d*_max_*, g*_age_) and for the biphasic VCBM are 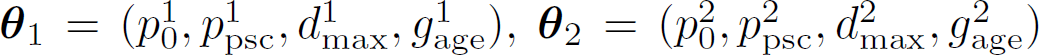 and *τ*. However, the complexity of the biological data simulating process may render the corresponding likelihood function computationally infeasible (Beaumont, 2019). When the likelihood function is intractable, standard Bayesian inference methods cannot be used to sample the posterior distribution. However, even when the likelihood function is intractable, it often remains feasible to generate simulations from the model.

#### 2.3.1 Approximate Bayesian Computation (ABC)

A method called approximate Bayesian computation (ABC) has been proposed to solve this problem by avoiding evaluation of the likelihood function (Sisson et al., 2018). The idea is to generate approximate samples from the parameter posterior by repeatedly simulating data ***x*** from the model for different sets of parameter values and assessing how ‘similar’ it is with the observed data ***y***. Parameter values that generate simulated data that are similar enough to the observed data are retained in the posterior sample. There are two main challenges in the successful implementation of ABC: choosing informative *summary statistics* (Prangle, 2015) and defining a suitable *distance function* (Drovandi and Frazier, 2022), *ρ*(***y*, *x***), that measures the ‘closeness’ of observed data with the simulated data. In our paper, the summary statistics are the observed data itself since the length of tumour growth time series data is short (i.e. *T* is small), and we select the distance function as

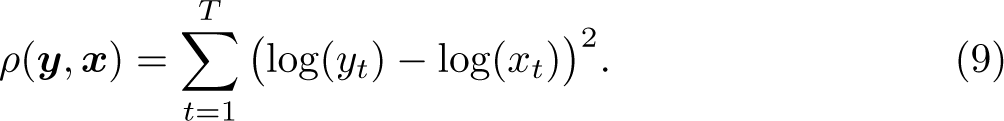

We compare the logarithm of the observed and simulated data since the tumour sizes tend to increase quickly with time. The approximate posterior implied by ABC is given by

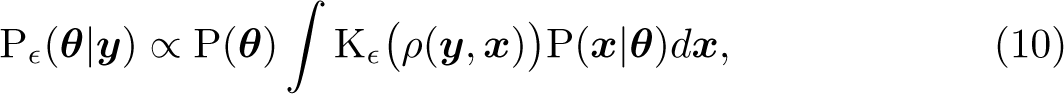

where K*_ɛ_* : ℝ_+_ 1→ ℝ_+_ is called a *kernel function* with *tolerance* parameter *ɛ*. Then, the approximate likelihood K*_ɛ_ ρ*(***y*, *x***) P(***x***|***θ***)*d****x*** can be estimated unbiasedly by using Monte Carlo via simulating ***x*** from the model. Here, the kernel function K*_ɛ_* is a weighting function that assigns higher weight to simulated data ***x*** that is closer to observed data ***y***. In this paper, we choose the indicator function as the kernel function, i.e., K*_ɛ_ ρ*(***y*, *x***) = I *ρ*(***y*, *x***) *< ɛ*.

Some popular algorithms like the ABC-rejection algorithm (Sisson et al., 2018; Warne et al., 2019) and Markov Chain Monte Carlo (MCMC)-ABC (Marjoram et al., 2003; Bortot et al., 2007; Sisson and Fan, 2011) have been used for parameter estimation in the context of biology (Beaumont, 2010; Csilĺery et al., 2010; Sunnåker et al., 2013; Beaumont, 2019). Since ABC rejection usually generates proposed values of ***θ*** from the prior, it can be inefficient if the target posterior differs substantially from the prior. The MCMC-ABC method is often computationally more efficient than ABC-rejection, since it aims to search locally around areas of high posterior probability density. However, MCMC-ABC can also be computationally costly, since the Markov chain can become stuck in low posterior regions, and, due to its serial nature, cannot easily exploit parallel computing architectures. On the other hand, the SMC-ABC algorithm can be more efficient since it can easily harness parallel computing, and it only proposes from the prior in the first iteration, and sequentially improves the proposal distribution. The SMC-ABC replenishment algorithm of Drovandi and Pettitt (2011) is used in a cell biology application in Carr et al. (2021), and we use the same algorithm to calibrate the VCBM. The SMC-ABC replenishment algorithm is explained in more detail in the next section.

#### 2.3.2 SMC-ABC replenishment algorithm

SMC-ABC samples from a sequence of increasingly accurate ABC posteriors based on defining a sequence of non-increasing tolerances *ɛ*_1_ ≥ *ɛ*_2_ ≥ · · · ≥ *ɛ_T_*:

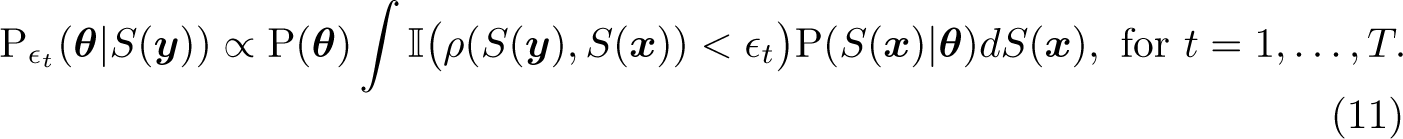

The algorithm first draws *N* independent samples from the prior P(***θ***), denoted here as 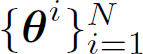. Then, for each draw of ***θ****^i^* (referred to as a particle), we simulate the stochastic model and compute its corresponding discrepancy *ρ^i^* to produce 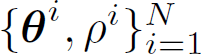. The particle set 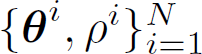 is then sorted by the discrepancy *ρ* such that *ρ*^1^ *< ρ*^2^ *<* · · · *< ρ^N^*. We then set the first tolerance threshold as *ɛ*_1_ = *ρ^N^*, i.e. the largest discrepancy value in the particle set. In order to propagate particles through the sequence of target distributions, the next tolerance is defined dynamically as *ɛ_t_* = *ρ^N−Na^* (where initially *t* = 2) where *N_a_* = ⌊*Na*⌋ and *a* is a tuning parameter that controls the adaptive selection of discrepancy thresholds and ⌊·⌋ is the floor function. Effectively, at each SMC iteration, we discard *a* × 100% of the particle set with the largest value of the discrepancy.

After discarding *a* × 100% of the particle set, there will only be *N* − *N_a_* particles remaining. In order to rejuvenate the particle set back to size *N*, we resample *N_a_* times with equal probability and replacement from the kept set of particles 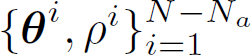, where both the parameter and discrepancy values are copied. Although the resampling step ensures that there are *N* particles in the set, it creates particle duplication.

In order to diversify the particle set, we apply an MCMC kernel to each of the resampled particles. The tuning parameters of the MCMC proposal distribution *q_t_*(·|·) can be obtained from the set of the particles following resampling, which are already distributed according to the ABC target with tolerance *ɛ_t_*. For example, if a multivariate normal random walk proposal is used, its covariance Σ*_t_* can be tuned based on the sample covariance computed from the particle set. Defining the current value of a particle we wish to move as ***θ***, we accept a proposal of parameter 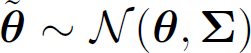 and simulated data 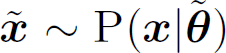 according to the probability

##### Algorithm 1

###### SMC-ABC replenishment algorithm (Drovandi and Pettitt, 2011)

**Input:** The observed data ***y***, the stochastic model P(***x***|***θ***), distance function *ρ*(·, ·), prior distribution P(***θ***), number of particles *N*, tuning parameters *a* and *c* for adaptive selection of discrepancy thresholds and selecting the number of MCMC iterations in the move steps, target tolerance *ɛ_T_*, initial number of trial MCMC iterations *S*_init_, minimum acceptable MCMC acceptance rate *p*_min_

**Output:** Samples, 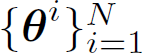, from the approximate posterior distribution

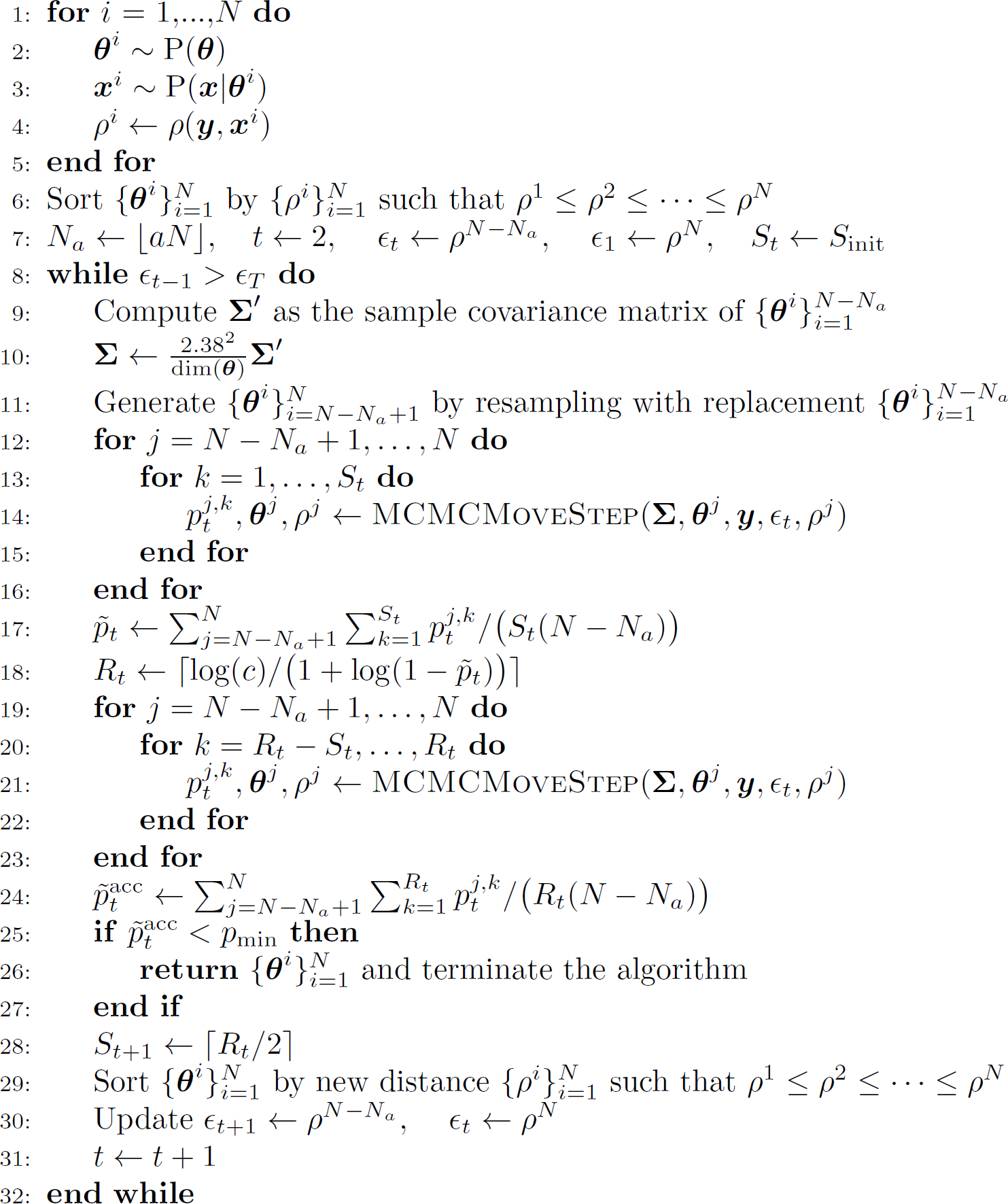

##### Algorithm 2

###### MCMC move step

**Input:** Covariance matrix **Σ**, parameter value ***θ***, observed data ***y***, tolerance *ɛ_t_* and discrepancy value *ρ*

**Output:** MCMC acceptance probability *p_t_*, parameter value ***θ*** and discrepancy value *ρ*

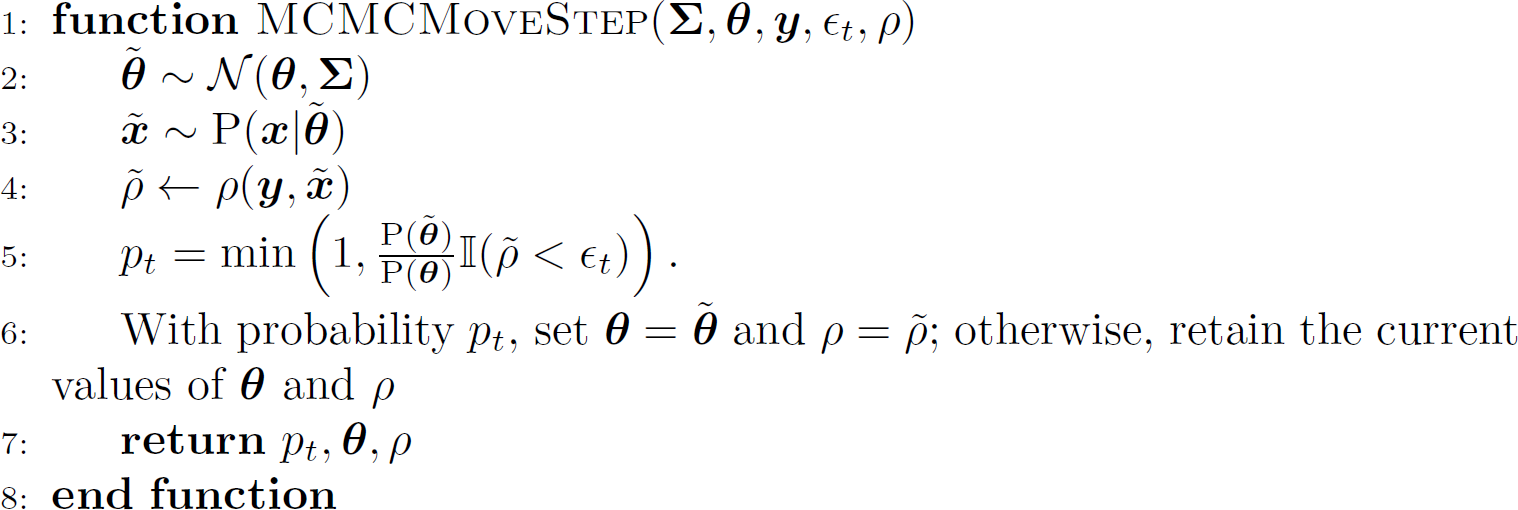

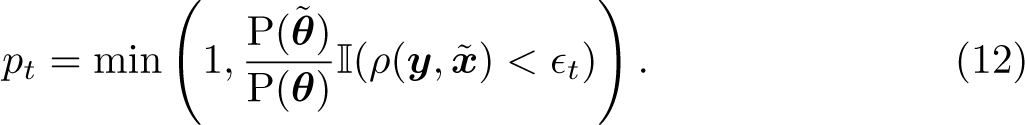

The proposal densities do not appear in the above acceptance probability due to the symmetry of the multivariate normal random walk proposal.

However, the above procedure may reject proposals and we may fail to move a large proportion of the resampled particles. Thus, we propose to apply *R_t_* iterations of the MCMC kernel to each resampled particle, so that there is

a high probability of moving each resampled particle at least once. Specifically, we set 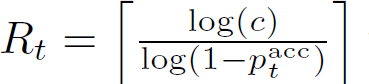 where *c* is a tuning parameter of the algorithm and is, theoretically, the probability that a particle is not moved in the *R_t_* iterations.

Here, 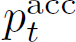 is the expected MCMC acceptance probability at the *t*th SMC iteration. Since 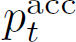 is unknown, we estimate it from *S_t_ < R_t_* trial MCMC iterations. Once it is estimated, we compute *R_t_* and perform the remaining *R_t_* − *S_t_* MCMC iterations on each of the resampled particles. For the next iteration we set the number of trial MCMC iterations to *S_t_*_+1_ = ⌈*R_t_/*2⌉.

There are two ways the algorithm can be stopped. One stopping rule is activated if some desired tolerance *ɛ_T_* is reached, i.e. the maximum discrepancy value in the particle set is below *ɛ_T_*. The second stopping rule is activated when the overall MCMC acceptance probability at a given SMC iteration falls below some user-defined *p*_acc_, implying that too much computation is required to progress the algorithm further. The overall acceptance rate can be estimated by

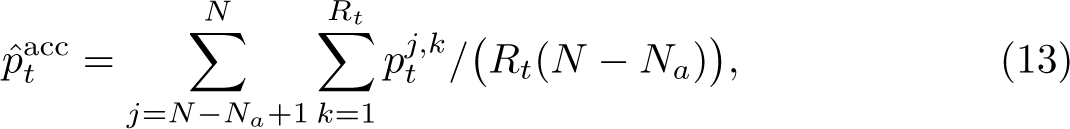

where 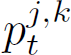 is the MCMC acceptance rate for particle *j* at MCMC iteration *k* and is obtained based on (12). For our study, we set the tuning parameters to be *a* = 0.5 and *c* = 0.01. We use the MCMC acceptance rate as the stopping rule of the algorithm and set *p*_min_ = 0.005.

### 2.4 Prior knowledge

The parameters *p*_0_ and *p*_psc_ are constrained to lie between zero and one as they are the *probabilities* of cell proliferation and cell invasion, respectively. Vague independent priors are used for each of these parameters, which, in each case, is a uniform distribution constrained by 0 and 1. For the switching time *τ*, we set its prior as a uniform distribution constrained by 2 and the time the maximum measurement taken. Since *τ* refers to the day that switching occuring, we take *τ* as integer. We assign a uniform distribution constrained by 0 and the maximum measurement day times 24 hours as prior to *g*_age_ so that the prior has a relatively large variance to create a reasonably vague prior.

Figure S1 depicts the distribution of distances *d* for the population of tumour cells based on the volume of the tumour. The *d* values obtained are proportional to the volume of the tumour, so as the tumour expands, more cells will be located further from its edge. It appears that *d* remains between 0 and 30 for ranges of tumour volumes relevant to the *in vivo* datasets. In other words, no cell has a distance greater than about *d* = 30. Hence, we assign a uniform distribution constrained by 0 and 50 as the prior for *d*_max_.

## 3 Results

In this section, we investigate the SMC-ABC algorithm’s ability to calibrate the VCBM. First, the SMC-ABC algorithm is applied to synthetic data generated from the model with known parameter values. Then, SMC-ABC is applied to three real tumour growth datasets to explore whether the VCBM can capture the tumour growth pattern for each type of cancer. The code and data is available via https://github.com/john-wang1015/Calibration BVCBM.

### 3.1 Validation with synthetic data

Prior to applying SMC-ABC to the observed experimental data sets, we first perform a preliminary investigation using synthetic datasets generated via simulation from the VCBM. This allows us to verify that the proposed SMC-ABC algorithm is able to produce expected results under simple settings, i.e. exhibit posterior concentration around the true values that were used to generate the data and for the posterior predictive distribution to be consistent with the generated data.

To validate the SMC-ABC method, we generate five sets of synthetic data using five “true” parameter settings (see Table S1 and plots for time series data in Fig S3). Three of the datasets are simulated with biphasic growth and two with monophasic growth, as the real data (see Fig 1) appear to exhibit both cases. For example, some mice in the pancreatic dataset appear to exhibit biphasic growth, whereas the mice in the breast cancer dataset appear to exhibit monophasic growth. The biphasic datasets use different values of *g*_age_ for the two phases. During the generation of simulated datasets, we initialize the tumour volume at 200*mm*^3^, consistent with the real data.

We use the posterior samples to approximate the posterior predictive distribution of the tumour volumes. In the supplementary document, we plot the (0.25, 0.75), (0.1, 0.9) and (0.025, 0.975) posterior predictive intervals. Our results (see Fig S5a, S6a, S7a, S8a and S9a) show that SMC-ABC can recover every synthetic dataset with reasonable accuracy, as the associated tumour volume falls within at least one of the intervals in the posterior predictive plots. The estimated univariate posterior distributions of synthetic time series (see Fig S5-S9) indicate that *g*_age_ is the largest driving force for tumour growth in the sense that the posterior is substantially more concentrated compared to the prior. However, the posteriors for *p*_0_ and *d*_max_ are not substantially different to the prior, and thus cannot be identified from the data.

Although Equation 6 has been widely used in ABM context to model tumour growth (Kansal et al., 2000a,b; Jiao and Torquato, 2011; Pourhasanzade et al., 2017; Jenner et al., 2020a), the sensitivity of the parameter *d*_max_ requires confirmation through simulation. Our simulation shows that *d*_max_ is only sensitive at a small scale, as seen in Fig S2a and S2b, but not for large values, such as 100 *mm*. This insensitivity may be due to the spring-based movement of cells, which allows new cells introduced through proliferation to have minimal impact on cells at the exterior. Consequently, the tumour edge and volume remain unaffected. As a result, the probability of cell division is not sensitive, as shown in Figure S1a.

### 3.2 Monotonic tumour growth data

In this section, we use SMC-ABC to estimate parameters of the monophasic VCBM from the ovarian and breast cancer datasets shown earlier in Figs 1a and 1b. We use only the monophasic model for these data sets, as we find that the standard VCBM provides a good fit to the data. Furthermore, the computational cost of fitting the standard VCBM with four parameters is substantially lower than fitting the biphasic VCBM with nine parameters. The parameter *g*_age_ appears to be the most sensitive parameter driving tumour growth in breast and ovarian datasets (see Figs 4 and 5). In the supplementary document, we show the posterior distributions and the bivariate plots for ovarian and breast tumour datasets from Figs S10-21.

**Fig. 4:**
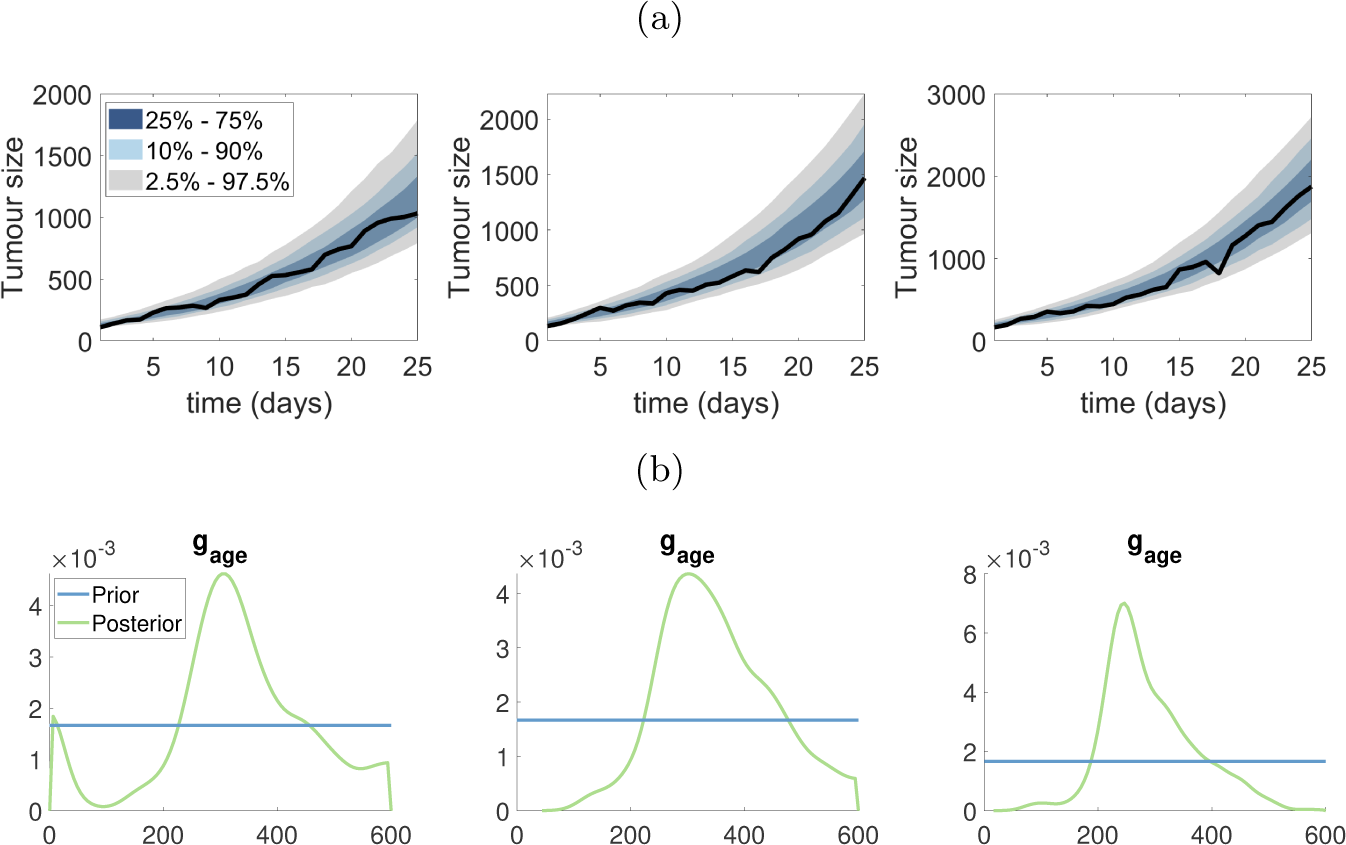
Estimating VCBM parameters for monophasic tumour growth fit using SMC-ABC to the ovarian cancer measurements. (a) The posterior predictive plots for each ovarian dataset; (b) The marginal distribution of *g*_age_ corresponding to the posterior predictive plots in (a).

**Fig. 5:**
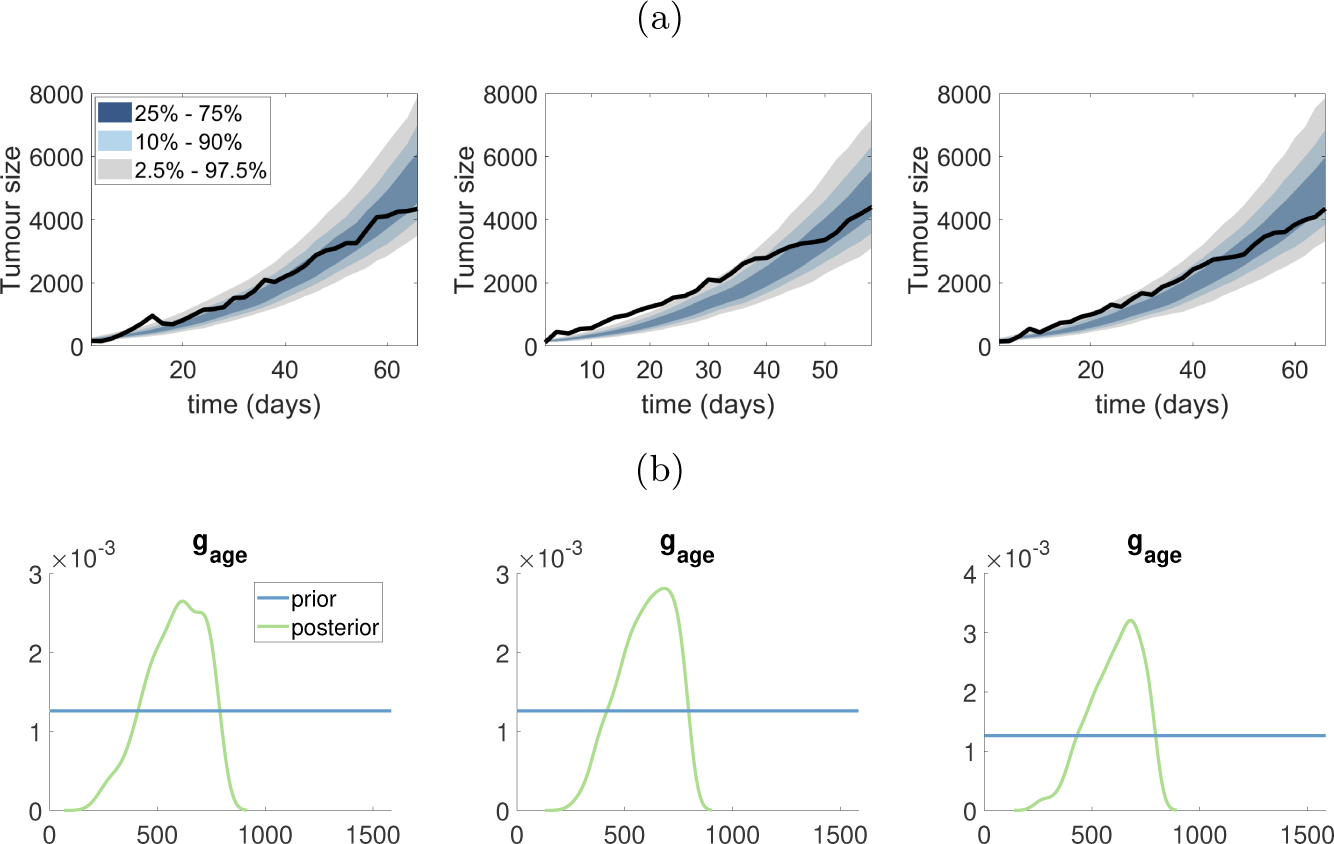
Estimating VCBM parameters for monophasic tumour growth fit using SMC-ABC to the breast cancer measurements. (a) The posterior predictive plots for each breast dataset; (b) The marginal distribution of *g*_age_ corresponding to the posterior predictive plots in (a).

The posterior predictive distributions for the ovarian and breast cancer datasets are shown in Fig 4a and Fig 5a, respectively. Firstly, it is evident that the VCBM provides a good fit to both cancer datasets, as the observed datasets lie comfortably within the prediction intervals. However, two of three datasets (first and third one) in the ovarian cancer dataset show a linear trend in volume size whereas the VCBM predicts exponential growth, and so the model is not as accurate at predicting the volume sizes at later times.

For both ovarian and breast cancer, the posterior distributions for *p*_psc_ and *d*_max_ are very similar to the prior, indicating that the data are not providing any additional information about these parameters. The posterior distribution for *p*_0_ is also similar to the prior for most datasets, except for some breast and ovarian datasets where there is some preference for smaller values of *p*_0_.

In contrast, the data are providing substantial information about *g*_age_, as indicated by a much more precise posterior distribution compared to the prior as shown in Figs 4b and 5b. This indicates that *g*_age_ strongly drives the dynamics of the VCBM, at least in terms of the tumour volume it produces. We also find there is positive correlation between *p*_psc_ and *g*_age_ in Fig S17, S19 and S21 in the ovarian cancer dataset. This indicates that as the probability of cell invasion increases, the growth rate of tumour cell will decrease to compensate.

### 3.3 Biphasic tumour growth data

In this section, SMC-ABC is employed to analyze the pancreatic cancer datasets, fitting the data with the biphasic VCBM. As observed in Fig. 1c, the first mouse in the pancreatic cancer dataset exhibits a change in growth pattern prior to day 15. Given this, we apply the biphasic VCBM to the entire pancreatic cancer dataset.

The posterior predictive distribution demonstrates that the biphasic VCBM offers a suitable fit for the four pancreatic datasets, as evidenced by the observed datasets falling within the prediction intervals as shown in the first column of Fig 6.

**Fig. 6:**
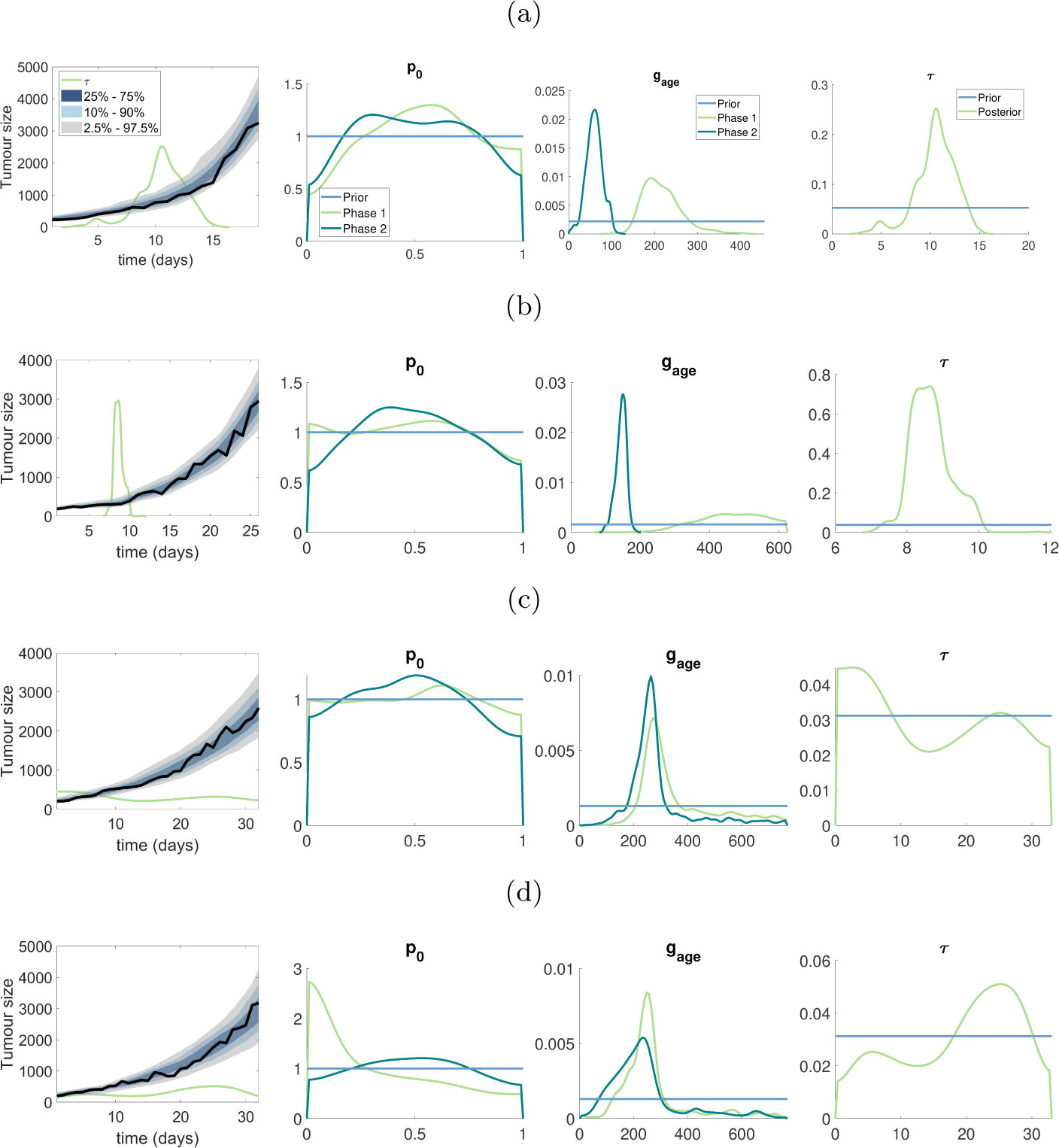
Results for pancreatic cancer dataset: (a)–(d) represent the results for the first to fourth mouse in pancreatic cancer dataset. The first plot in (a)–(d) shows the posterior predictive distribution, the green line shows the scaled posterior distribution for *τ* used to indicate the switching time of tumour growth, the true density for posterior *τ* is in final plot of (a)–(d). The second and third plots for (a)–(d) show the marginal posterior for *p*_0_ and *g*_age_ in two phases. The horizontal blue lines represent the prior distribution.

Our analysis of pancreatic tumor growth using the biphasic VCBM reveals that *g*_age_ is an important parameter describing the tumor growth dynamics. Changes in the *g*_age_ parameter can reflect alterations in the tumor microenvironment and genetic or epigenetic alterations in the tumor cells. For the first two mice in the pancreatic dataset, there is a significant difference in *g*_age_ between the two phases of growth (as exhibited by non-overlapping posterior distributions). Furthermore, for these two mice, the switching time between the two growth phases is reasonably well identified. For the third and fourth mice in the pancreatic dataset, the posterior distribution of *g*_age_ between the two growth phases has significant overlap, and the posterior distribution of the switching time is close to uniform over the experiment. This suggests that these two mice exhibit monophasic growth, and demonstrates how the biphasic VCBM can effectively reduce to a standard VCBM. In the supplementary document, we present the bivariate plots for parameters in Figs S22 - S25. A follow-up analysis could then apply the standard VCBM for these two mice, as the extra complexity of the biphasic VCBM is not warranted for these data sets.

## 4 Discussion

In this research, we develop a biphasic off-lattice ABM based on a Voronoi tessellation, which is an extension of Jenner et al. (2020a). We demonstrate the utility of the new model and the standard VCBM by calibrating them to real world tumour growth time series data.

The Voronoi Cell-Based model (VCBM) describes the stochastic nature of cancer cell proliferation and has an intractable likelihood function making it challenging to fit to data. The SMC-ABC replenishment algorithm suggested by Drovandi and Pettitt (2011) is suitable for calibrating the VCBM since the likelihood of the VCBM is intractable. While the SMC-ABC replenishment technique has been widely employed Carr et al. (2021); Varghese et al. (2020); Vo et al. (2015); Warne et al. (2020), this is one of the first times that an ABM like this has been calibrated to actual tumour growth data. The outcome of our calibration not only verifies the conclusion of Jenner et al. (2020a) that *g*_age_ is the most sensitive parameter, but also demonstrates the biphasic VCBM can capture the switching of tumour growth dynamics and precisely estimate the value of *g*_age_ in different phases.

In our study, we use breast cancer, pancreatic cancer and ovarian cancer tumour growth measurements *in vivo* to examine the robustness and flexibility of the VCBM. It is evident that *d*_max_ and *p*_psc_ are not informed by any of the data sets. This is most likely due to the fact that tumour volume is most impacted by the expansion of the tumour periphery, which only requires the cells on the periphery to proliferate. As such, modelling the probability of proliferation as a function of the distance to the tumour periphery is not informed by simply tumour volume measurements and requires more informative data collection. In contrast, *g*_age_ provides rich information about the average time a cell takes to proliferate.

It is evident that some of the pancreatic datasets appear to have biphasic tumour growth patterns. To investigate this, we assumed there was some time at which tumour growth could vary and introduce *τ* which is a parameter that control the time at which the tumour growth dynamics change. Performing SMC-ABC, we saw that *d*_max_ and *p*_psc_ are not able to provide any information as the posterior for both phases are close to the prior. However, the *g*_age_ posteriors indicate clearly that the average time for cell proliferation is different based on the two phases. Where it is clear that in the first phase, the average time to proliferation is slower than in the second phase. This highlights a shift in tumour growth which may relate to the expansion of the tumour in the subcutaneous tissue or potentially a mutation in the cell cycling. However, to conclude the cause of this shift needs more investigation from both an experimental and modelling point of view.

To reduce computational cost, we implemented the VCBM in only 2-dimensional space. In this way, we approximate tumour growth *in vivo* in 2-dimensions which we feel matched the experimental measurements which are also only in two dimensions. However, having the third dimension measurement of the tumour *in vivo* would provide us with an understanding of the irregularity in tumour volume that we may be missing by only considering 2-dimensional tumour growth. In future work, we hope to use 3-dimensional tumour volume measurements, such as Magnetic resonance imaging (MRI), to calibrate the 3-dimensional form of the VCBM and improve model accuracy. This would allow us to recapitulate tumour shape irregularities, which we currently assume are negligible in the 2-dimensional VCBM.

As our results suggest, the pancreatic cancer data set suggests that biphasic tumour growth can occur. The next steps for our model are to reformulate the cellular proliferation to be a function of tumour space, so that as the tumour grows we may be able to capture multiphasic growth. One idea for this, would be to consider sub-clonal populations within the tumour that arise stochastically with varying proliferation constants.

While the SMC-ABC is a relatively fast off-line algorithm, it is still computationally expensive as for every proposed parameter value generated in a Bayesian algorithm, a full dataset must be simulated. When simulation time is non-trivial, this then creates a computationally intensive calibration algorithm. On the other hand, online algorithms can iteratively update model parameter estimates as data are introduced sequentially. Such an approach would be useful for our application, since we would only need to simulate data forward one time step at each iteration and future work hopes to investigate this further.

Overall, given the slow adoption of likelihood-free algorithms that can infer parameters in ABMs, we feel our manuscript provides inspiration for others using ABMs applied in a biological context where data is available to attempt parameter inference. In turn, our results suggest that not all parameters are practically identifiable in the VCBM, which is information only gained through attempting to infer these parameters to data. Since the underlying structure of the VCBM is used in many different applications, this finding has a flow-on effect for currently published models, whose parameters may not be identifiable. It also motivates future experiments that may be used to identify these parameters, such as time-series flow cytometry measurements to identify cell proliferation markers. Lastly, the clear ability of our algorithm to identify the biphasic switching time of *in vivo* tumour growth suggests biologically that tumours grown subcutaneously in mice may in reality exhibit two phases. This then allows us the understand more deeply how *in vivo* tumour growth may differ from in *situ* tumour growth in humans and help inform our understanding of experimental findings.

In conclusion, we have applied SMC-ABC to calibrate the standard VCBM and biphasic extension with real breast cancer, pancreatic cancer and ovarian cancer tumour growth datasets. It is evident that *g*_age_ is the informative parameter for the VCBM and allows it to recapitulate *in vivo* tumour growth data. Unfortunately, *p*_psc_ and *d*_max_ are non-identifiable. We also find the biphasic VCBM shows the ability to extract information from biphasic tumour growth datasets like a pancreatic tumour. We intend to extend the VCBM so that it can capture potentially multiphasic tumour growth pattern and also improve the computational cost of SMC-ABC by moving the algorithm online.

## Acknowledgement

We thank the computational resources provided by QUT’s High Performance Computing and Research Support Group (HPC). We also thank the authors of the previously published experimental data Dr Chae-Ok Yun, Dr Kara Perrow and Dr Samantha Wade for supplying the original published data sets. Xiaoyu Wang and Christopher Drovandi were supported by an Australian Research Council Future Fellowship (FT210100260). Adrianne L. Jenner was supported by the QUT Early Career Researcher Scheme. The project was partly supported by the Centre for Data Science First Byte grant.

## Statements and Declarations

The authors have no competing interests to declare that are relevant to the content of this article.

