## supplementary document for "Calibration of Agent Based Models for Monophasic and Biphasic Tumour Growth using Approximate Bayesian Computation"

### List of Tables

S1 Parameters used in the generation of the three synthetic datasets. . . . . . 5

### 1 Analysis of $d_{\max}$

(a)

(b)

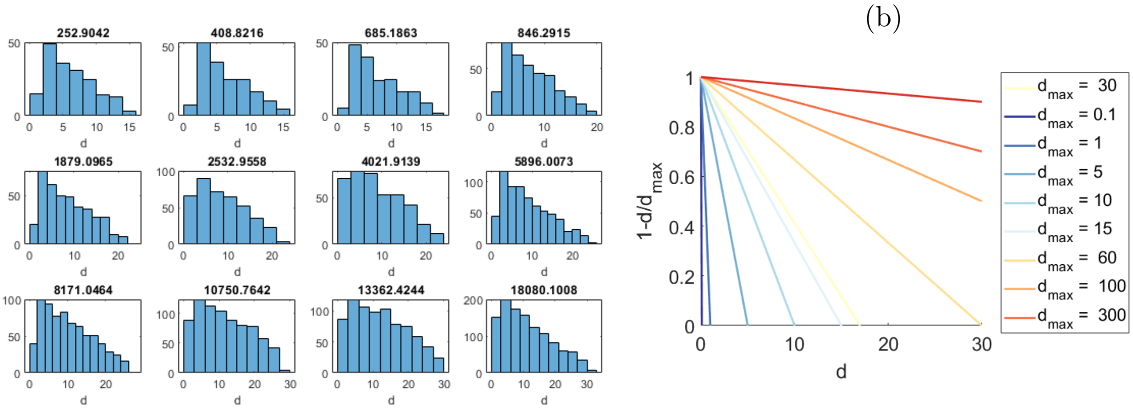

Figure S1: The distribution of distance $d$ for the population of tumour cells based on the volume of the tumour. (a) is the histograms of $d$ in different number of cells; (b) shows the relationship between $d$ and $1 - d/d_{\max}$.

(a)

(b)

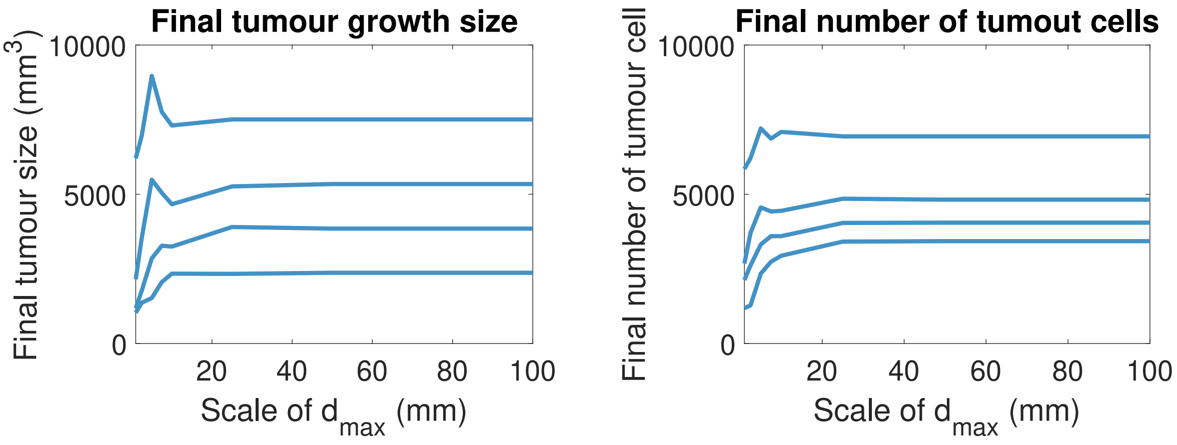

Figure S2: The effect of different scale of $d_{\max}$ in VCBM while other model parameters fixed: (a) final tumour data generated from model for different scale of $d_{\max}$; (b) final total number of cells generated from model for different scale of $d_{\max}$.

#### 2 Synthetic dataset

Table S1: Parameters used in the generation of the three synthetic datasets.

| Dataset | $p_0$ | $p_{\mathrm{psc}}$ | $d_{\max}$ | $(g_{\mathrm{age}_1}, g_{\mathrm{age}_2})$ | $\tau$ | length (days) |
|---|---|---|---|---|---|---|
| 1 | 1 | 0 | 17.3 | (300, 100) | 16 | 32 |
| 2 | 1 | 0 | 17.3 | (200,75) | 16 | 32 |
| 3 | 1 | 0 | 17.3 | (100,300) | 16 | 32 |
| 4 | 0.4 | $10^{-5}$ | 31 | 155 | N/A | 25 |
| 5 | 0.9 | $10^{-5}$ | 20 | 171 | N/A | 66 |

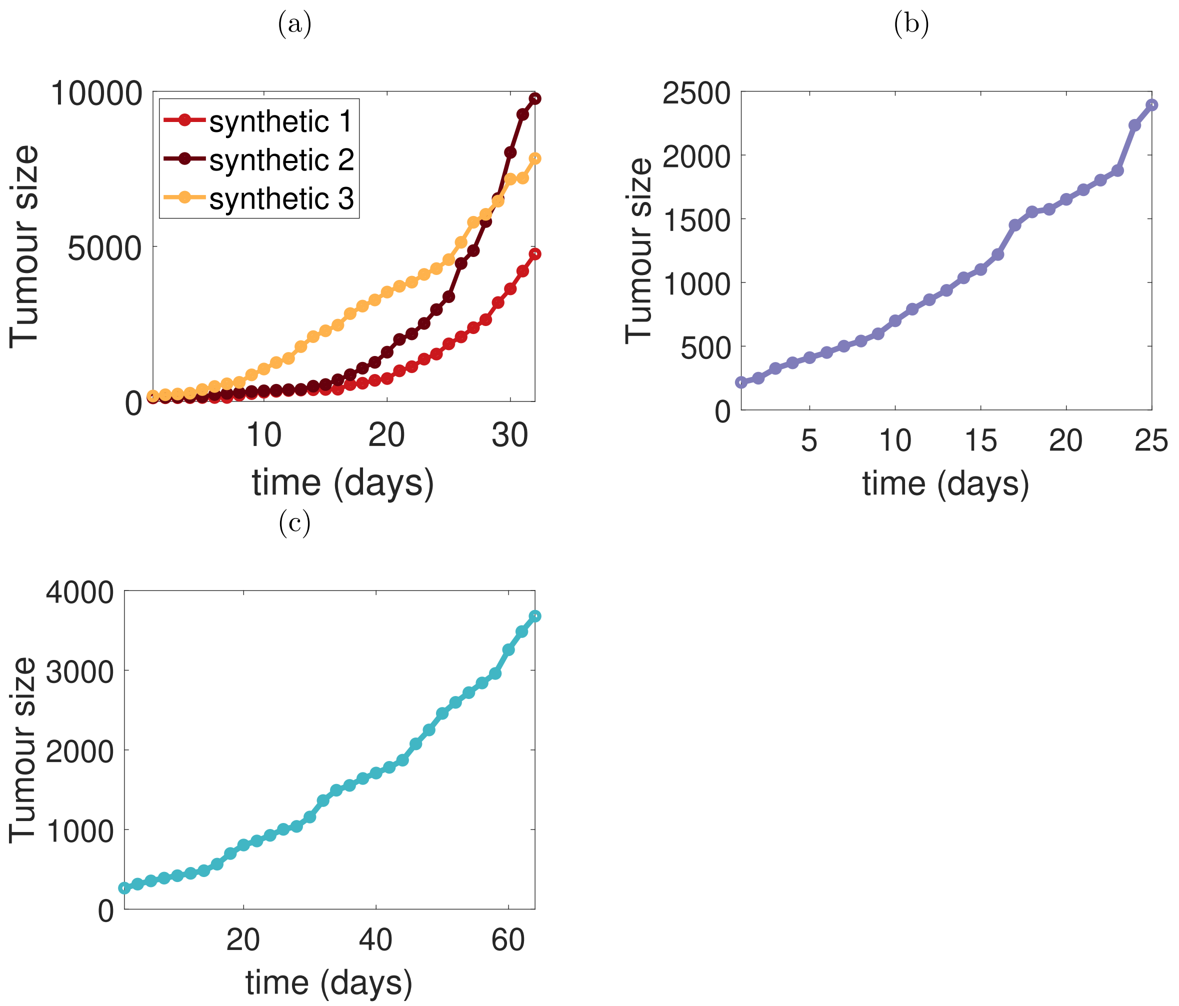

Figure S3: Synthetic tumour volume measurements. (a) is the synthetic time series dataset 1 to 3; (b) is the synthetic time series dataset 4 and (c) is the synthetic time series dataset 5.

### Prior Predictive distributions

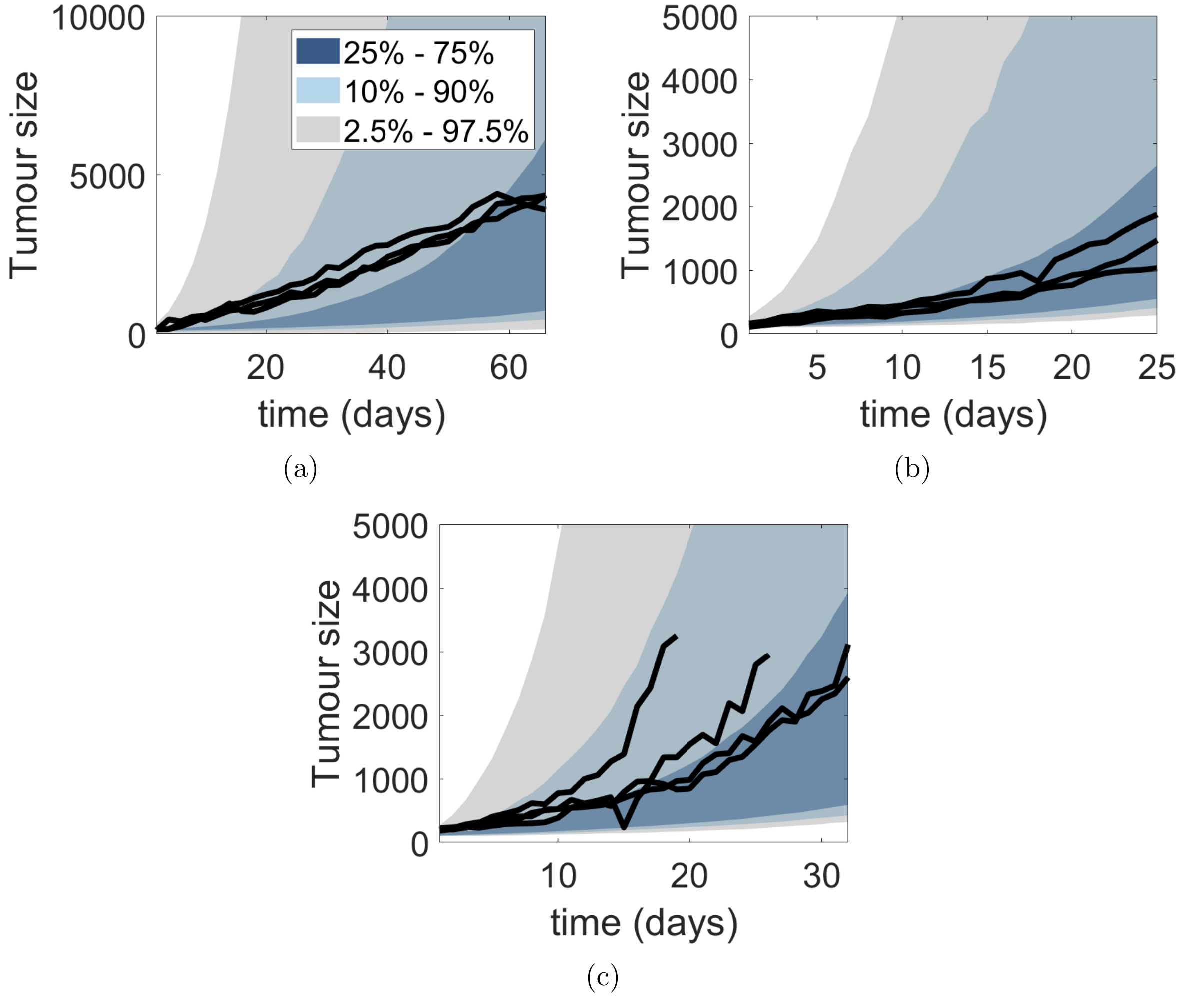

Figure S4: Prior predictive distributions for each of experimental datasets. Black solid lines in (a) - (c) are breast, ovarian and pancreatic tumour datasets, respectively.

### 3 Posterior for synthetic datasets

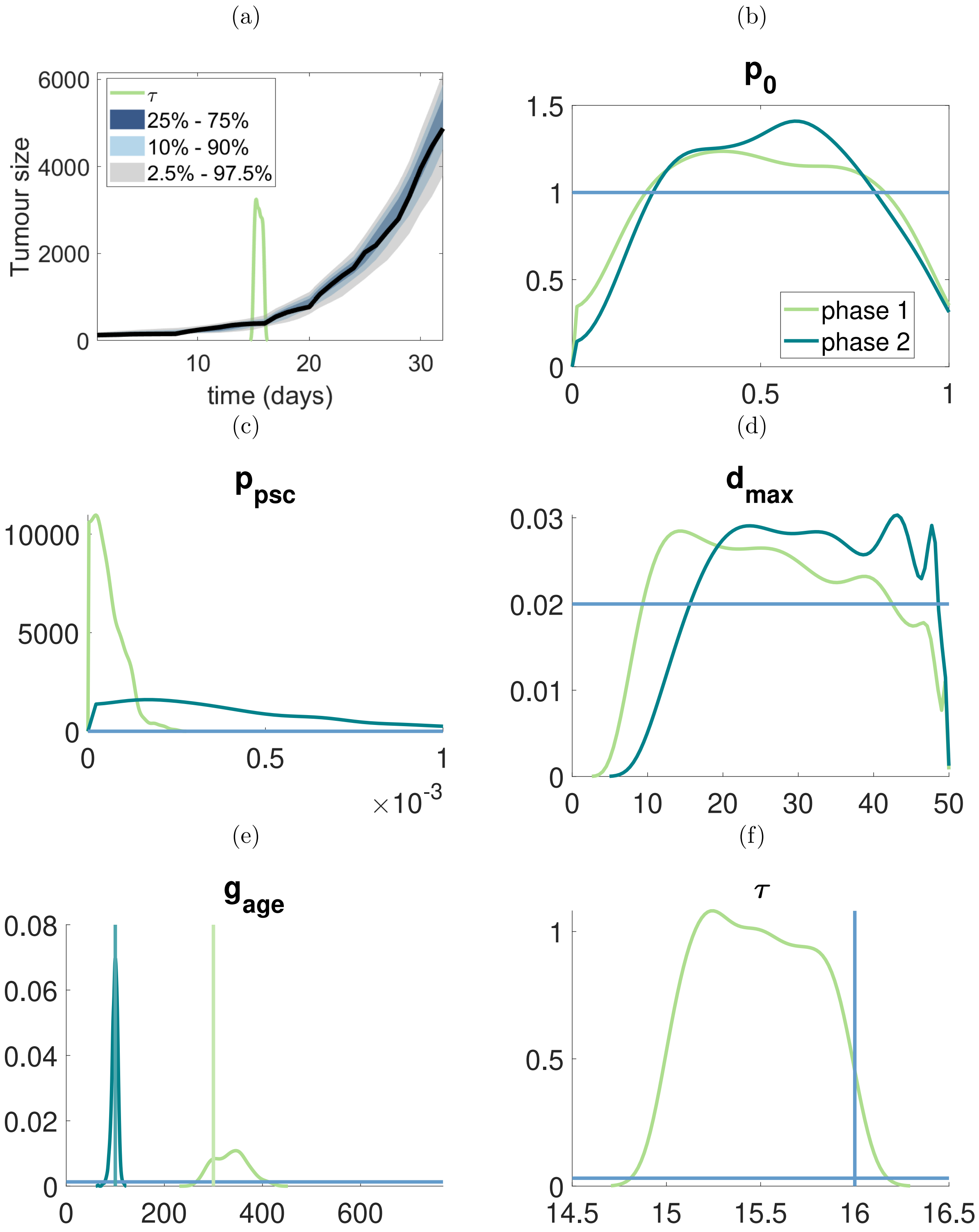

Figure S5: Results for synthetic dataset 1: (a) shows the posterior predictive distribution, the green line shows the scale up posterior distribution for $\tau$ used to indicate the switching time of tumour growth, the true density for posterior $\tau$ is in (f); (b) - (f) show the marginal posterior for each of parameter, the horizontal blue lines represent the prior distribution and the vertical line in (e) and (f) refers to the "true" values of parameters.

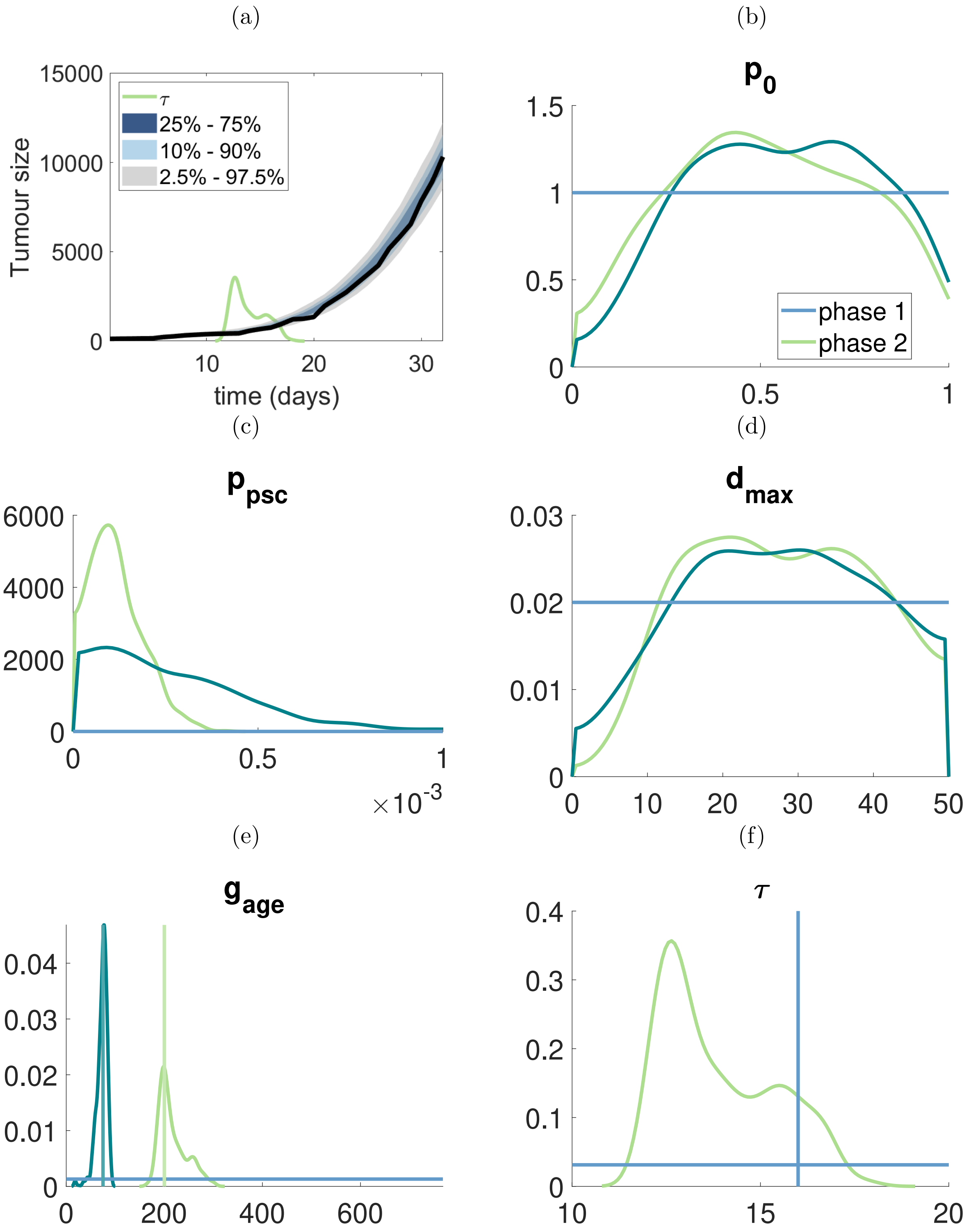

Figure S6: Results for synthetic dataset 2: (a) shows the posterior predictive distribution, the green line shows the scale up posterior distribution for $\tau$ used to indicate the switching time of tumour growth, the true density for posterior $\tau$ is in (f); (b) - (f) show the marginal posterior for each of parameter, the horizontal blue lines represent the prior distribution and the vertical line in (e) and (f) refers to the "true" values of parameters.

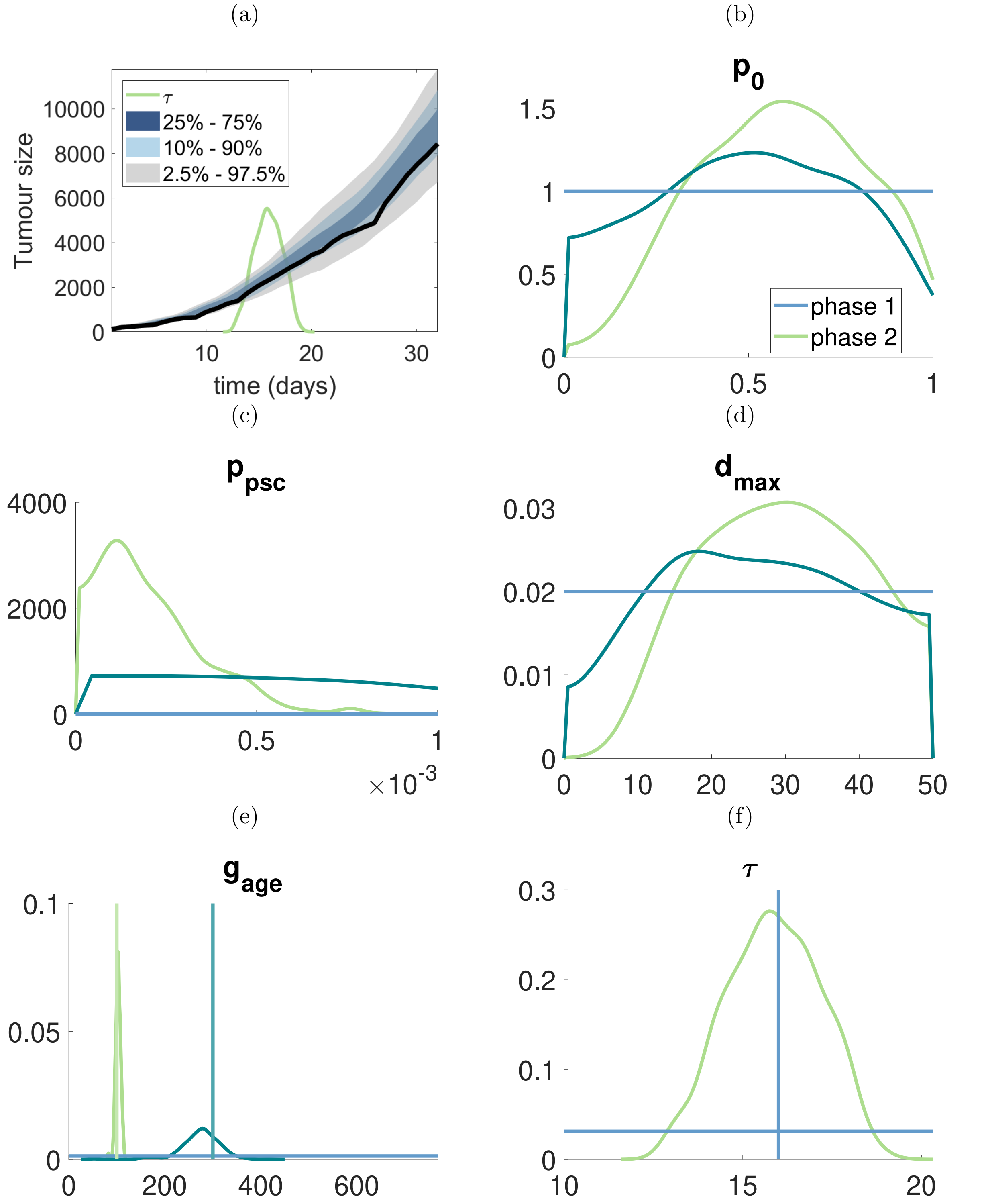

Figure S7: Results for synthetic dataset 3: (a) shows the posterior predictive distribution, the green line shows the scale up posterior distribution for $\tau$ used to indicate the switching time of tumour growth, the true density for posterior $\tau$ is in (f); (b) - (f) show the marginal posterior for each of parameter, the horizontal blue lines represent the prior distribution and the vertical line in (e) and (f) refers to the "true" values of parameters.

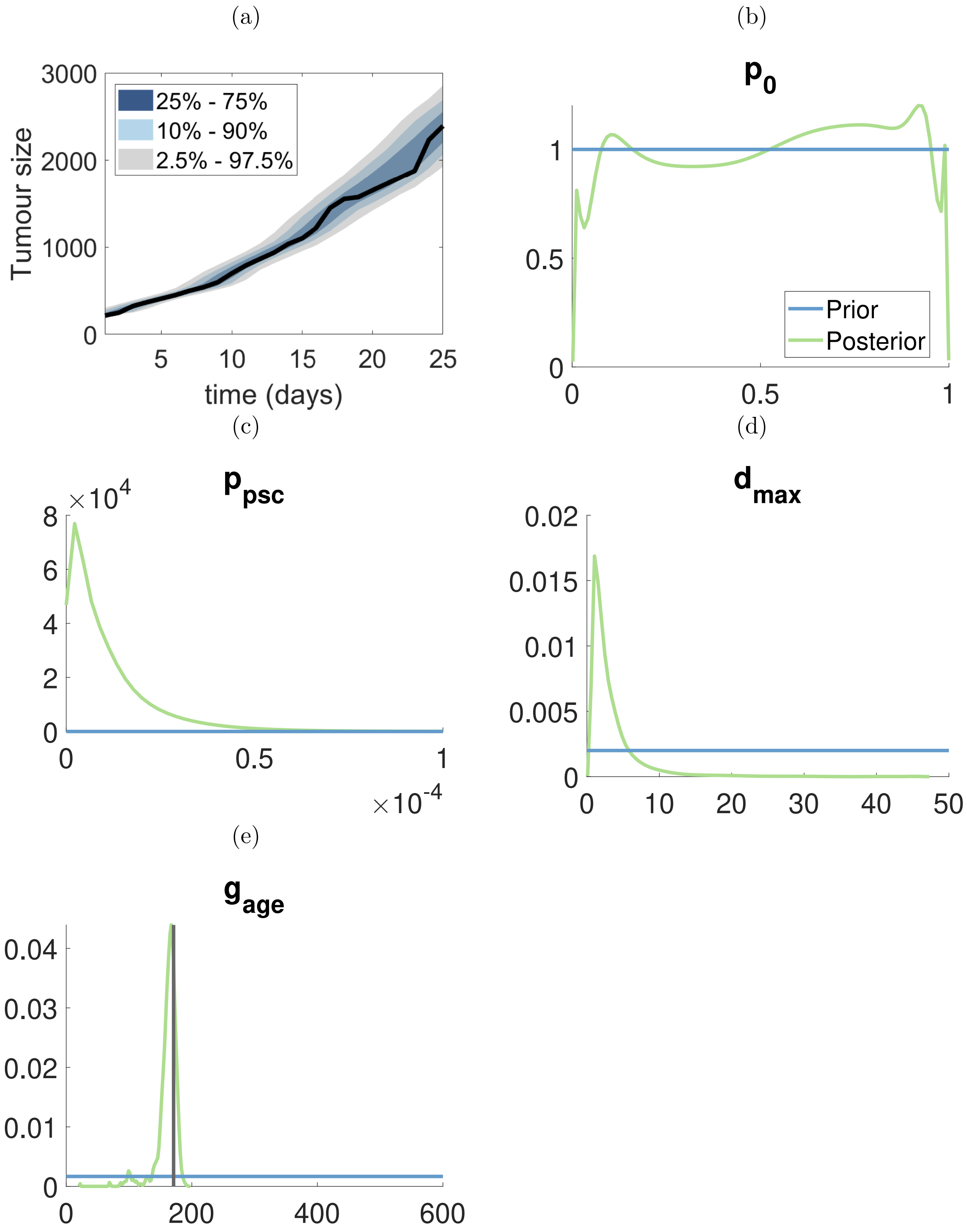

Figure S8: Results for synthetic dataset 4: (a) shows the posterior predictive distribution; (b) - (e) show the marginal posterior for each of parameter, the horizontal blue lines represent the prior distribution and the vertical line in (e) refers to the “true” values of parameters.

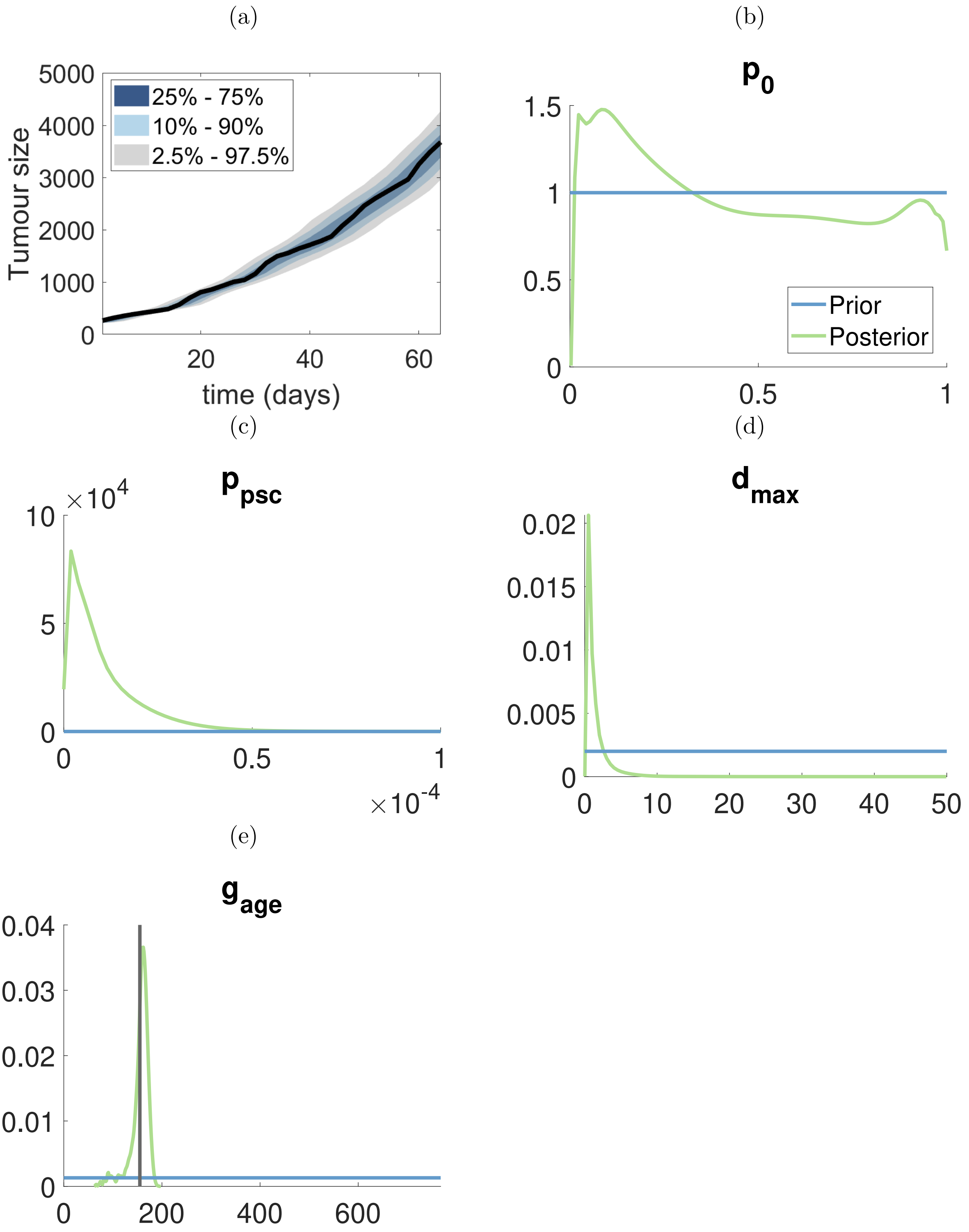

Figure S9: Results for synthetic dataset 5: (a) shows the posterior predictive distribution; (b) - (e) show the marginal posterior for each of parameter, the horizontal blue lines represent the prior distribution and the vertical line in (e) refers to the “true” values of parameters.

### 4 Posterior for breast tumour datasets

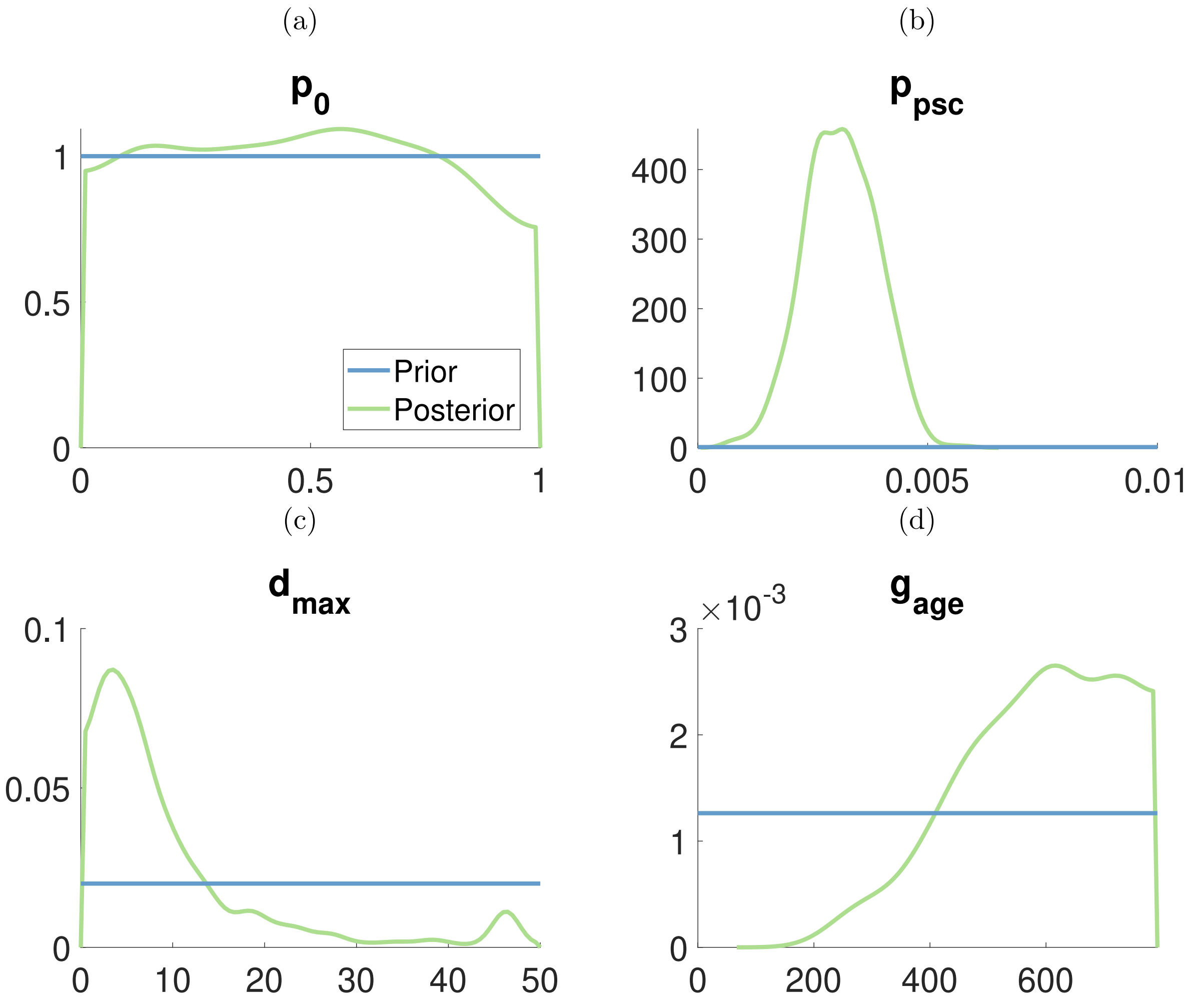

Figure S10: Marginal posterior distributions for first mouse in breast cancer dataset. The horizontal blue lines represent the prior distribution and green lines represent to marginal posterior distributions.

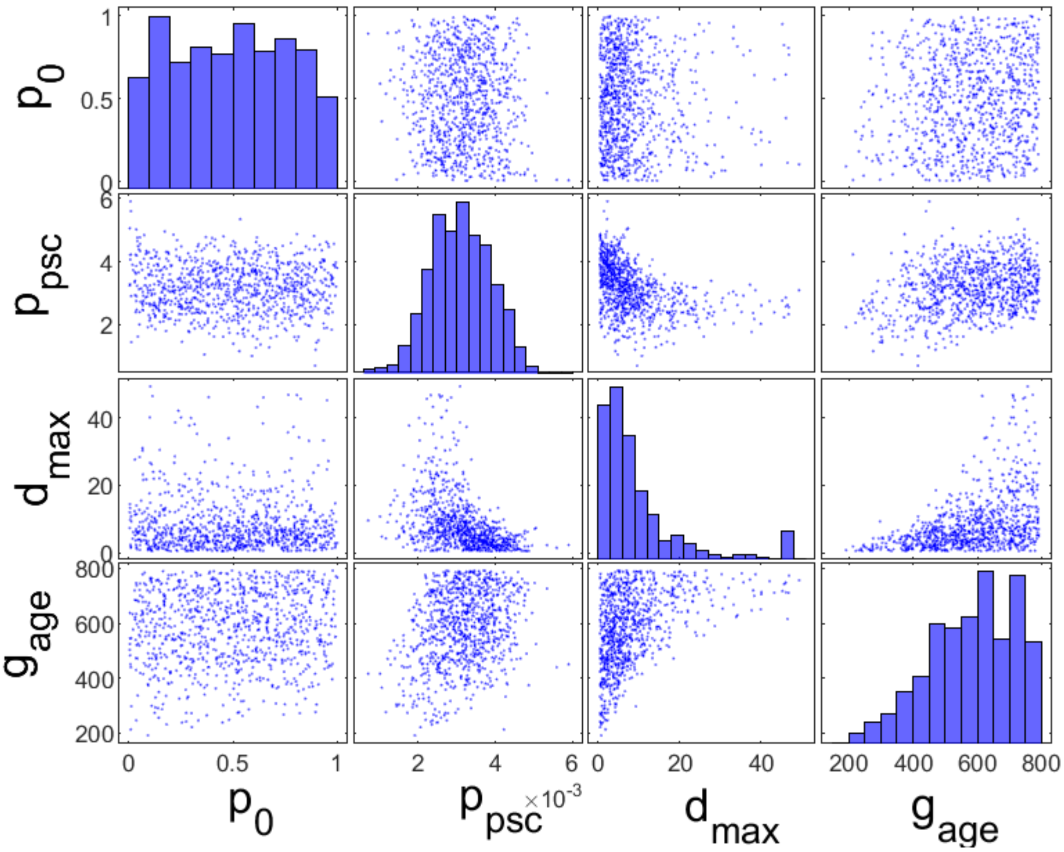

Figure S11: Bivariate plot for first mouse in breast cancer dataset.

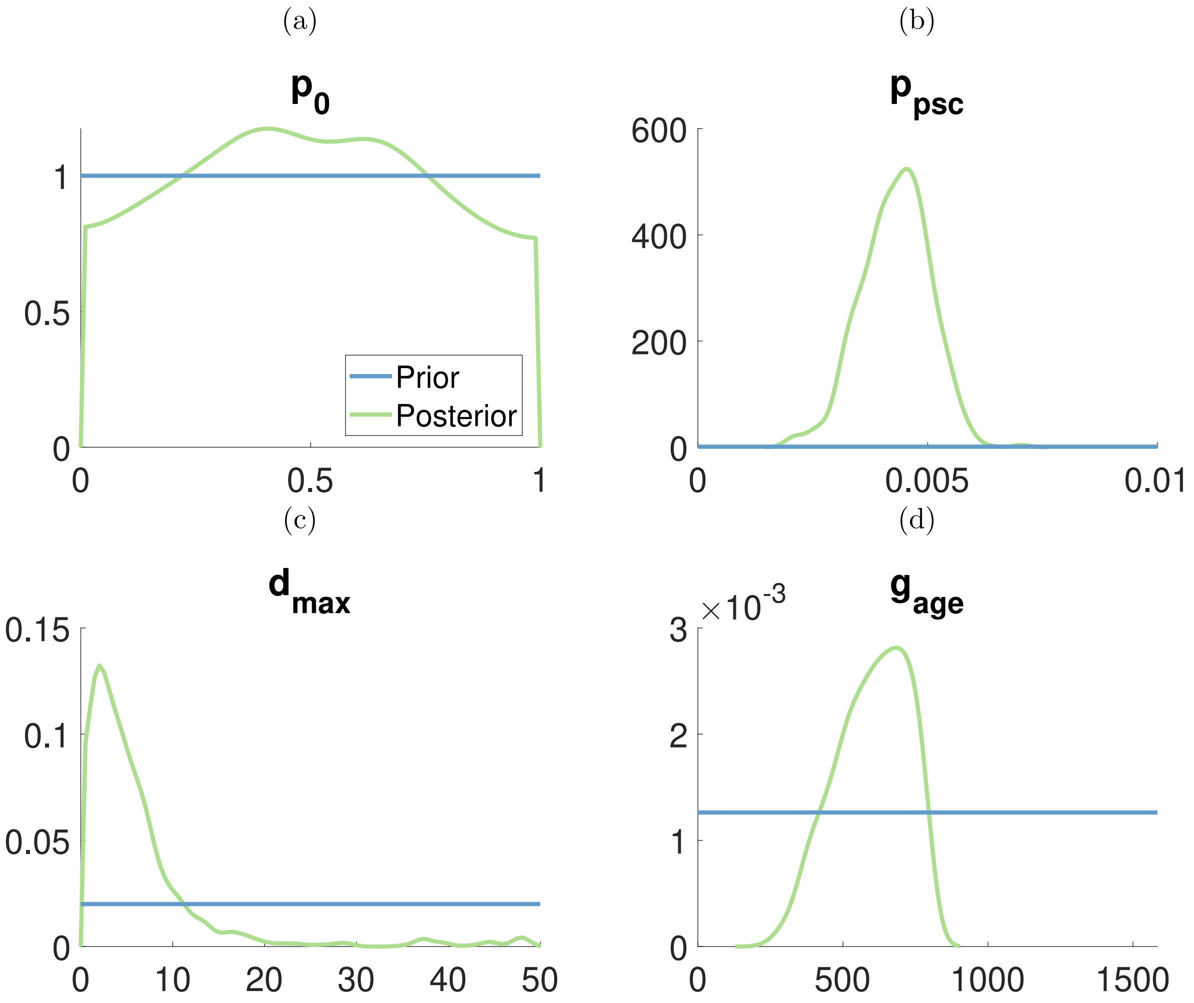

Figure S12: Marginal posterior distributions for second mouse in breast cancer dataset. The horizontal blue lines represent the prior distribution and green lines represent to marginal posterior distributions.

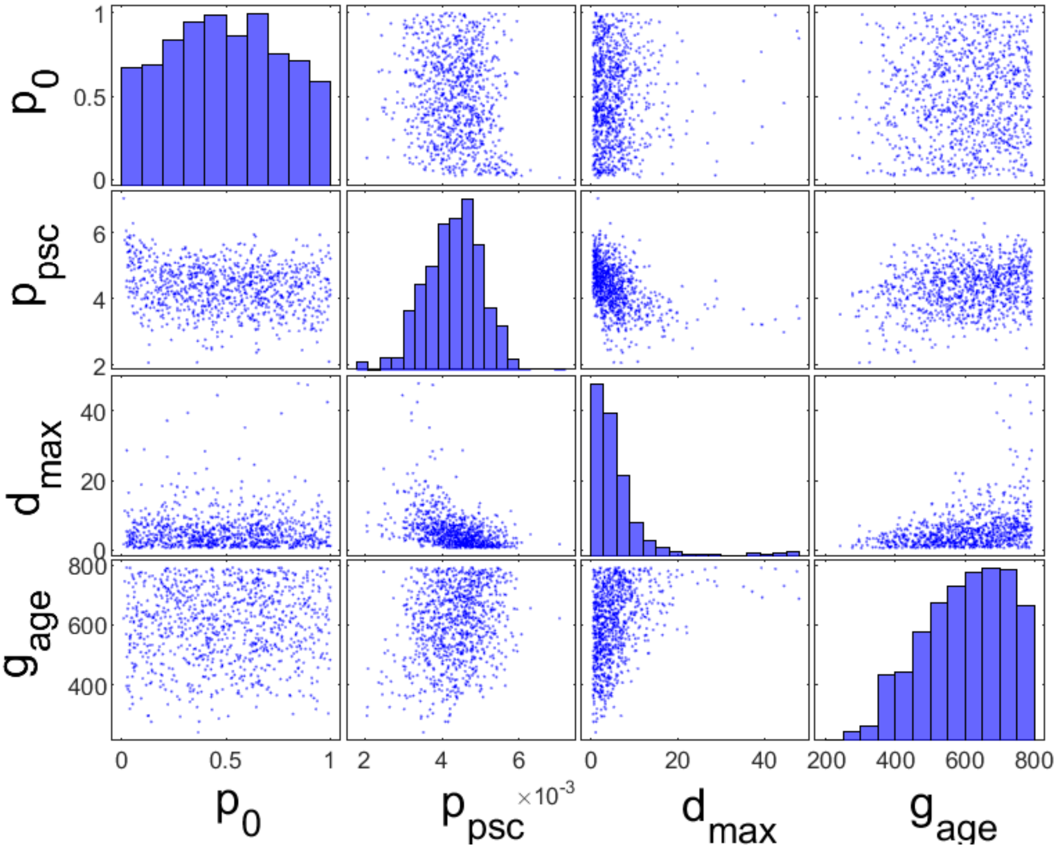

Figure S13: Bivariate plot for second mouse in breast cancer dataset.

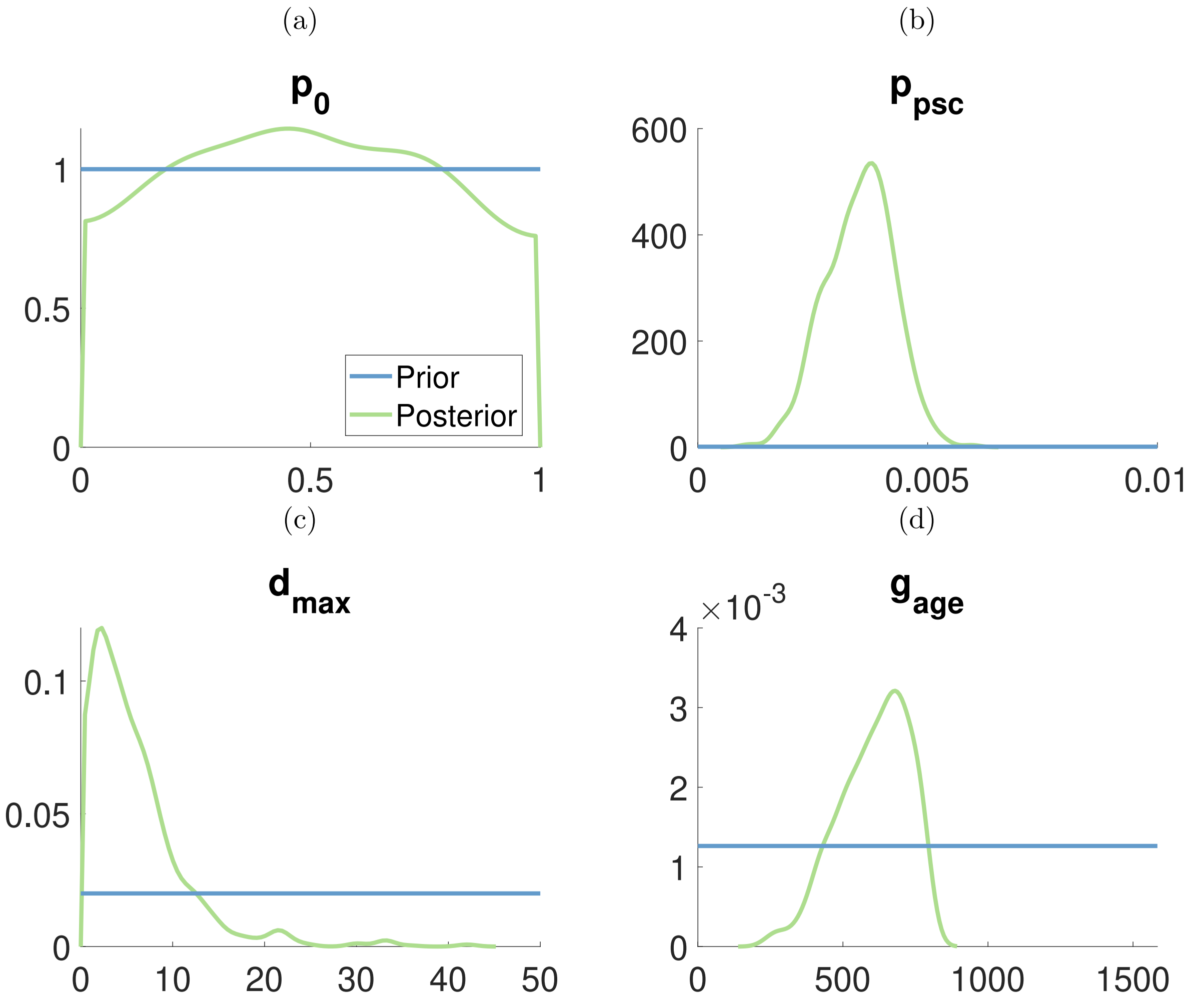

Figure S14: Marginal posterior distributions for third mouse in breast cancer dataset. The horizontal blue lines represent the prior distribution and green lines represent to marginal posterior distributions.

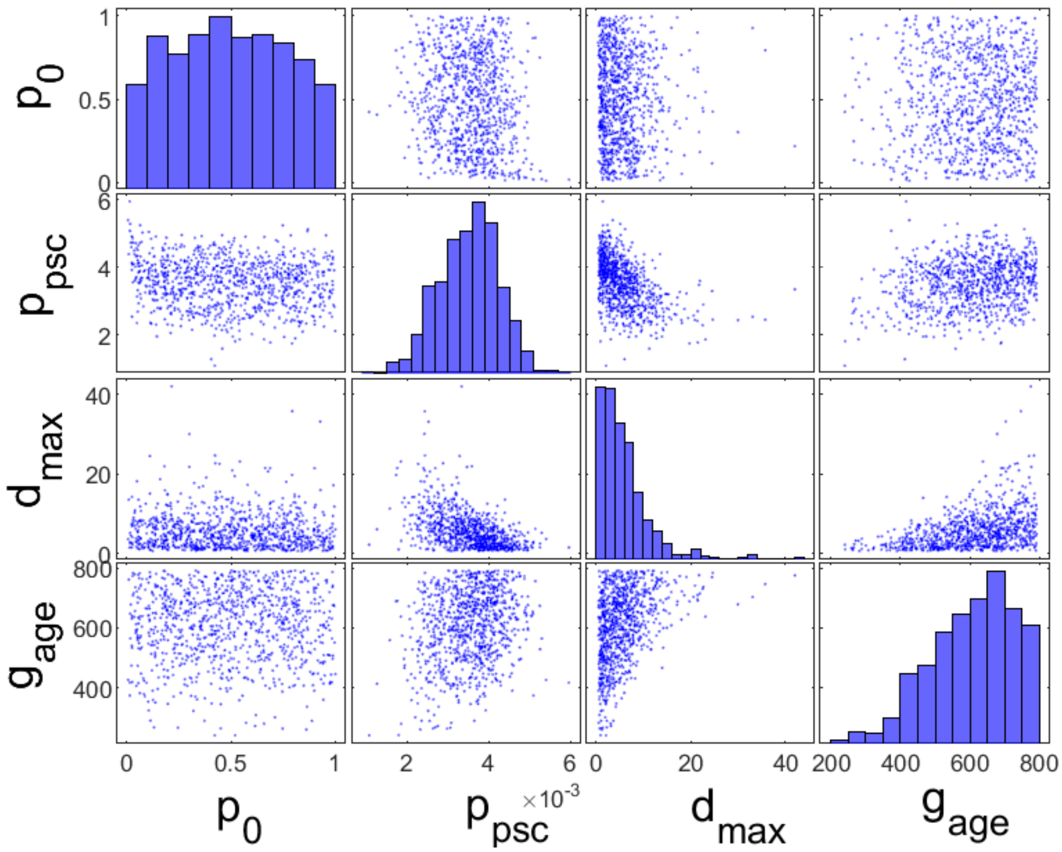

Figure S15: Bivariate plot for third mouse in breast cancer dataset.

### 5 Posterior for ovarian tumour datasets

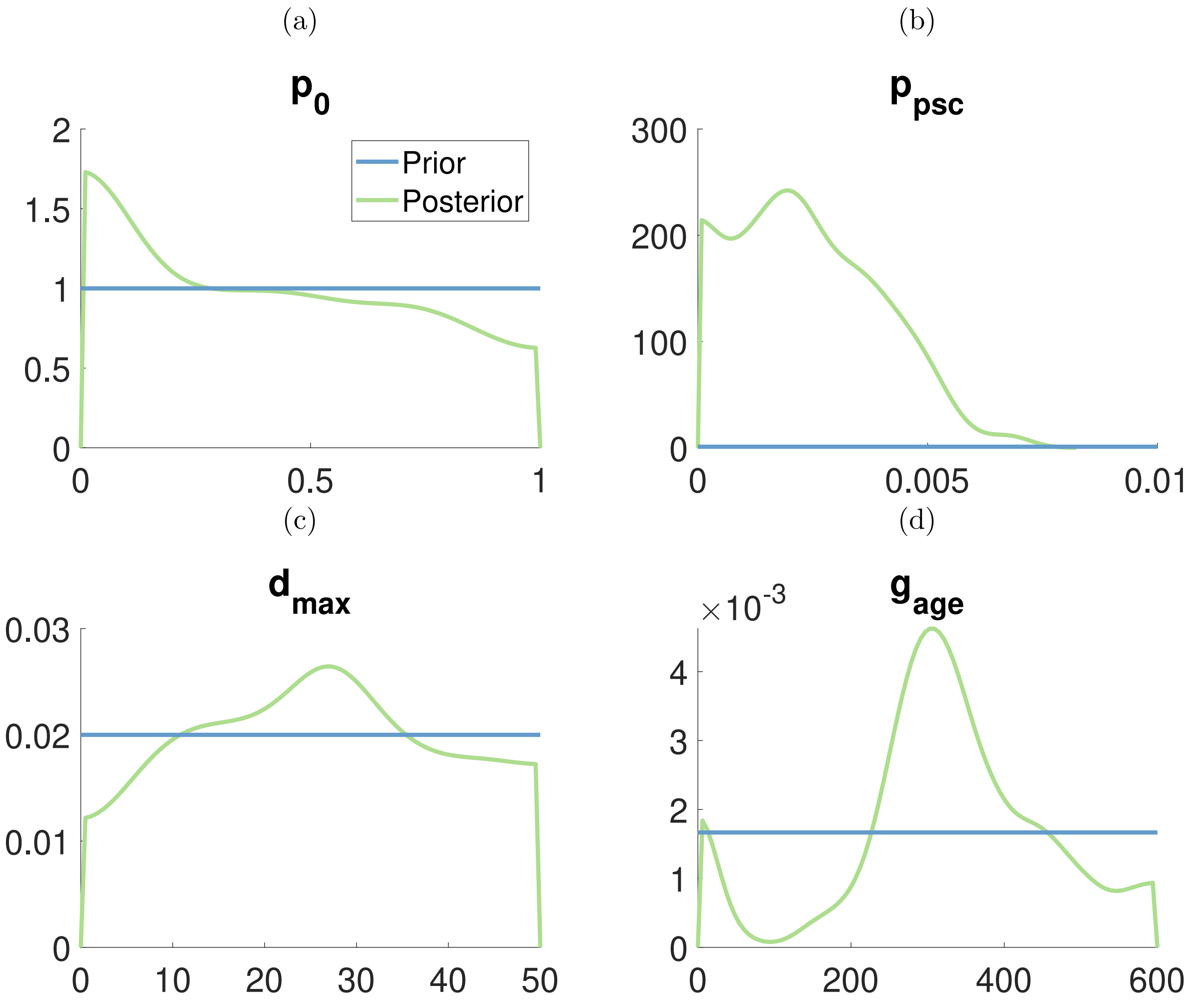

Figure S16: Marginal posterior distributions for first mouse in ovarian cancer dataset. The horizontal blue lines represent the prior distribution and green lines represent to marginal posterior distributions.

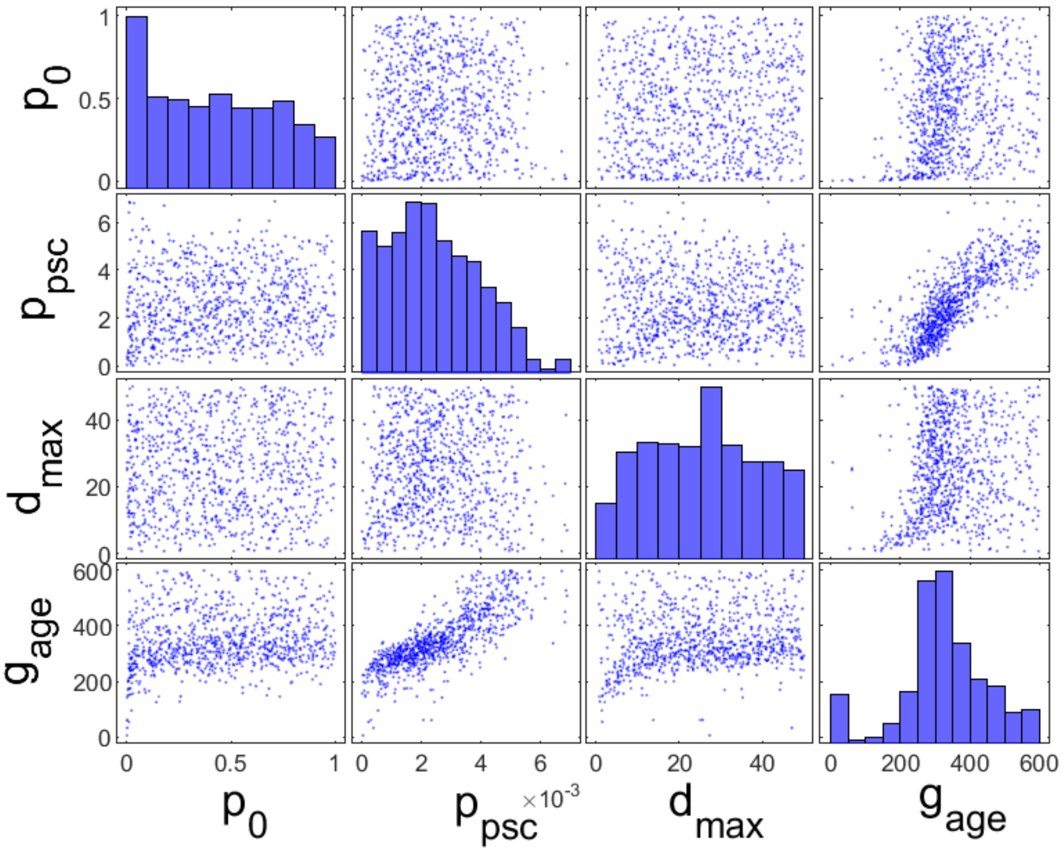

Figure S17: Bivariate plot for first mouse in ovarian cancer dataset.

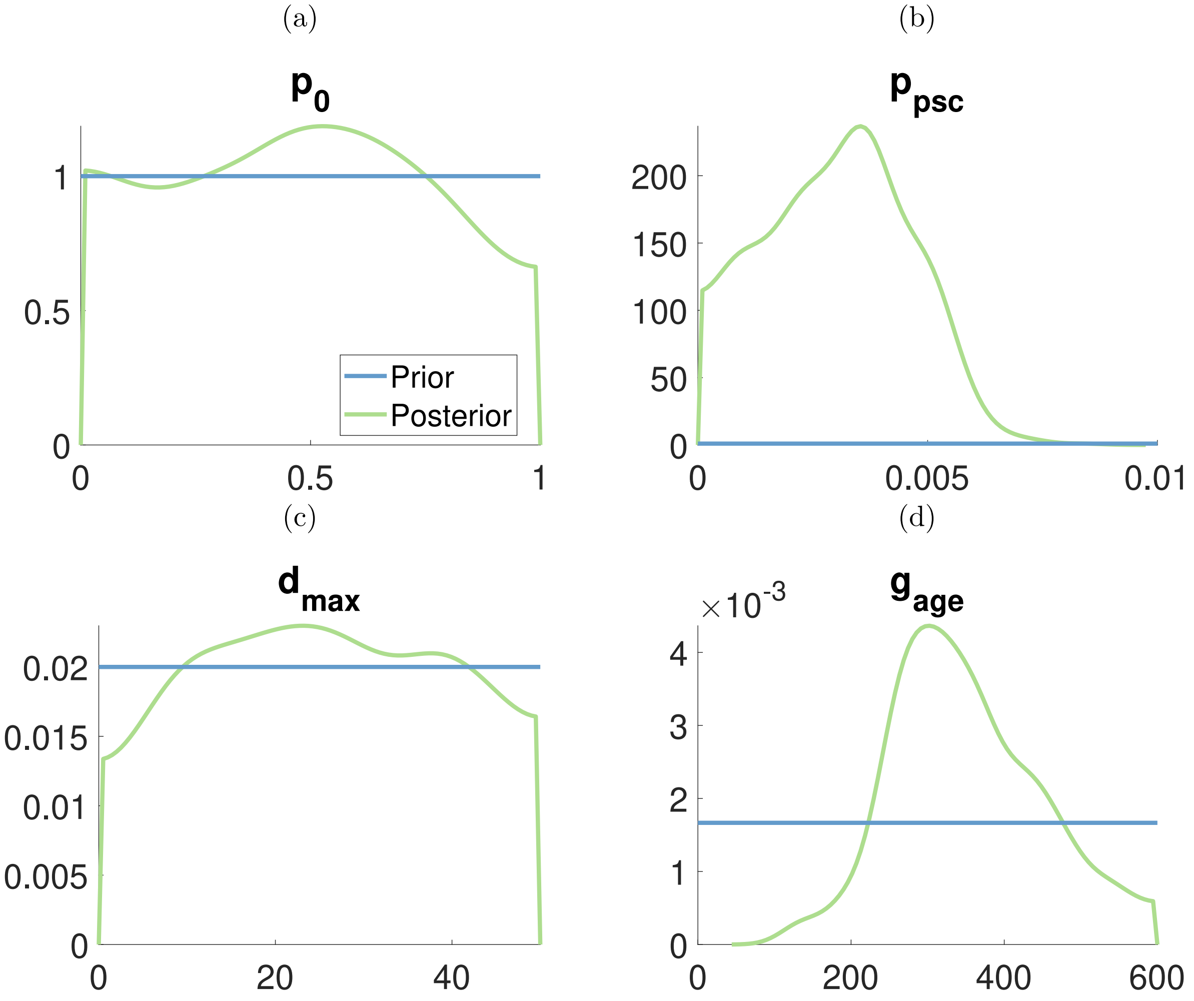

Figure S18: Marginal posterior distributions for second mouse in ovarian cancer dataset. The horizontal blue lines represent the prior distribution and green lines represent to marginal posterior distributions.

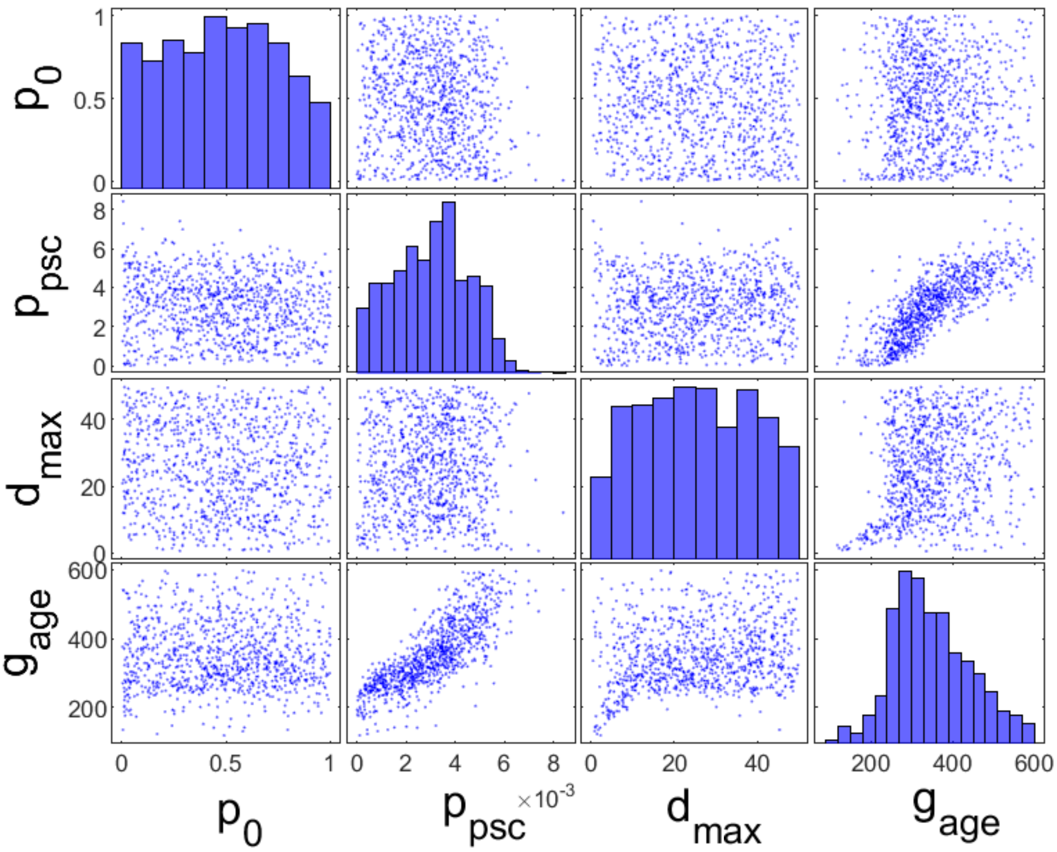

Figure S19: Bivariate plot for second mouse in ovarian cancer dataset.

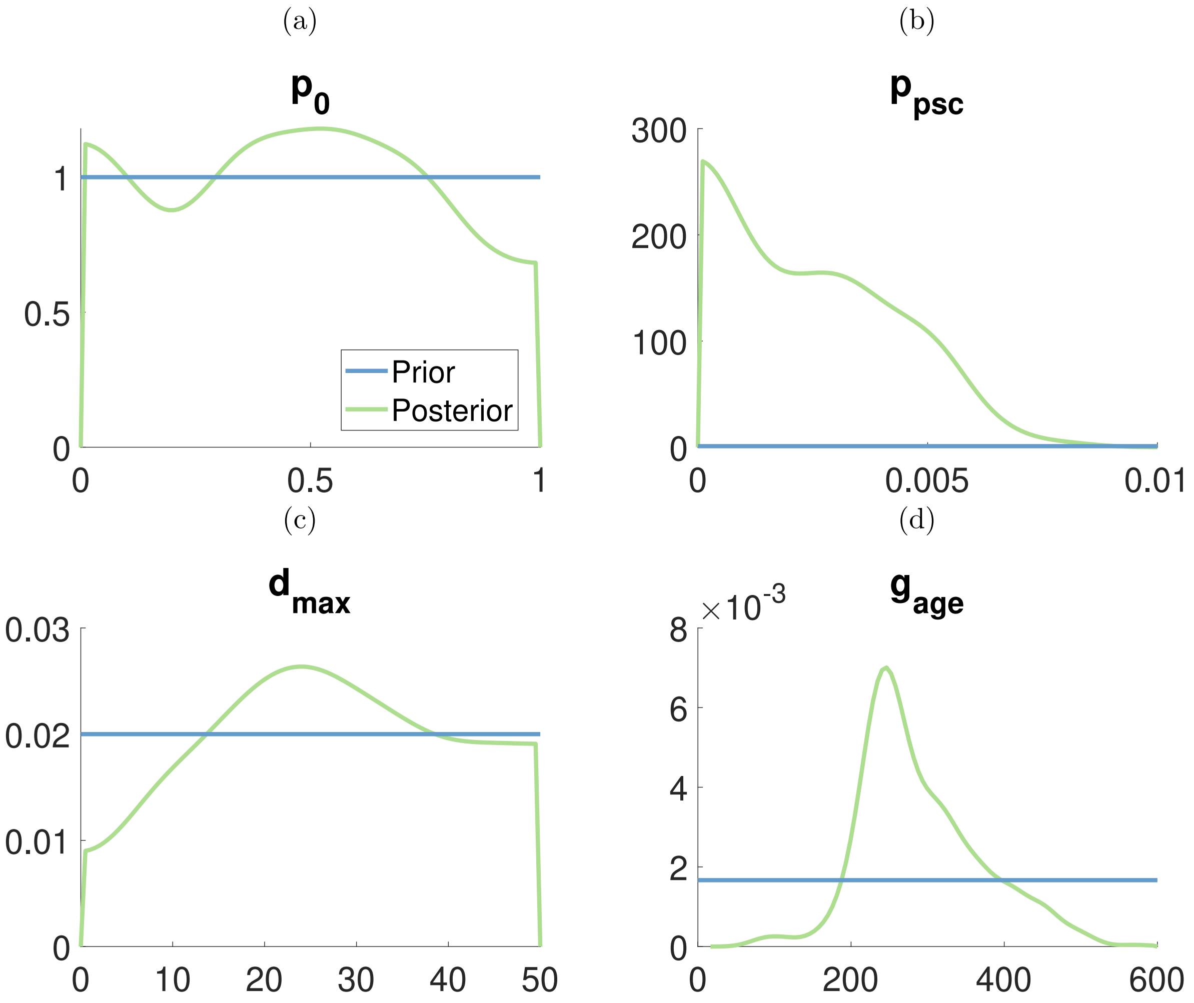

Figure S20: Marginal posterior distributions for third mouse in ovarian cancer dataset. The horizontal blue lines represent the prior distribution and green lines represent to marginal posterior distributions.

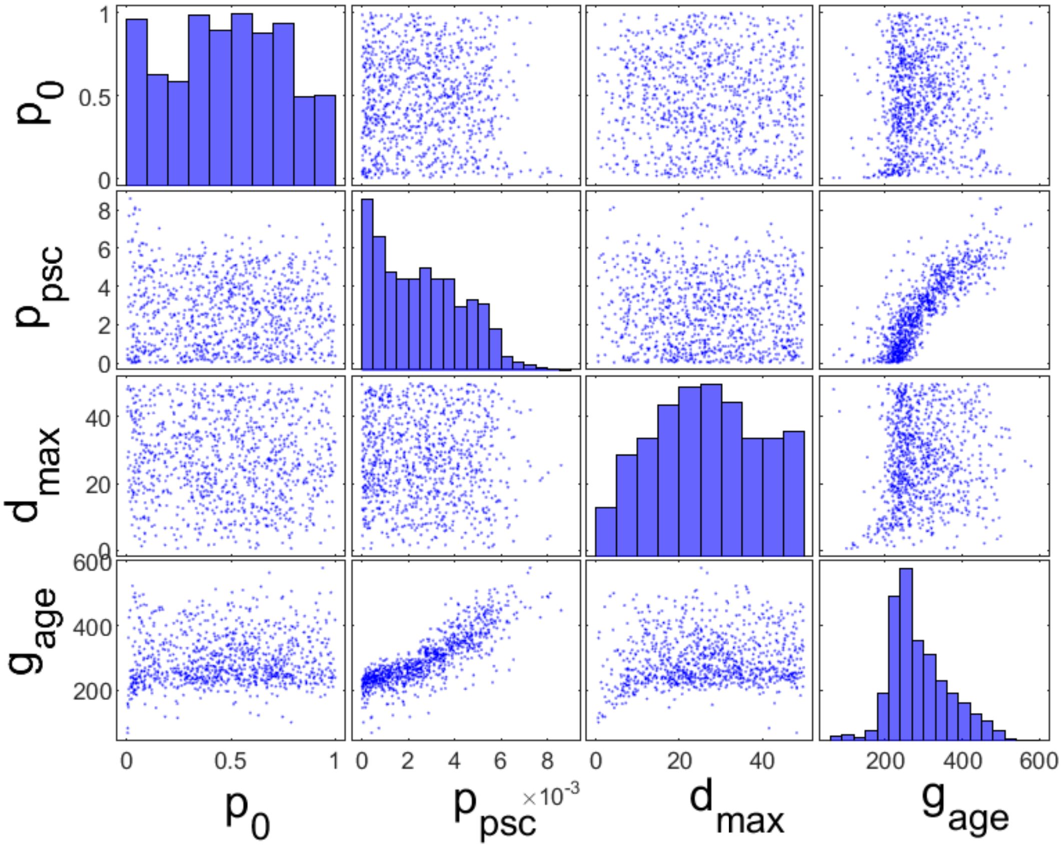

Figure S21: Bivariate plot for third mouse in ovarian cancer dataset.

### 6 Bivariate plots for pancreatic tumour datasets

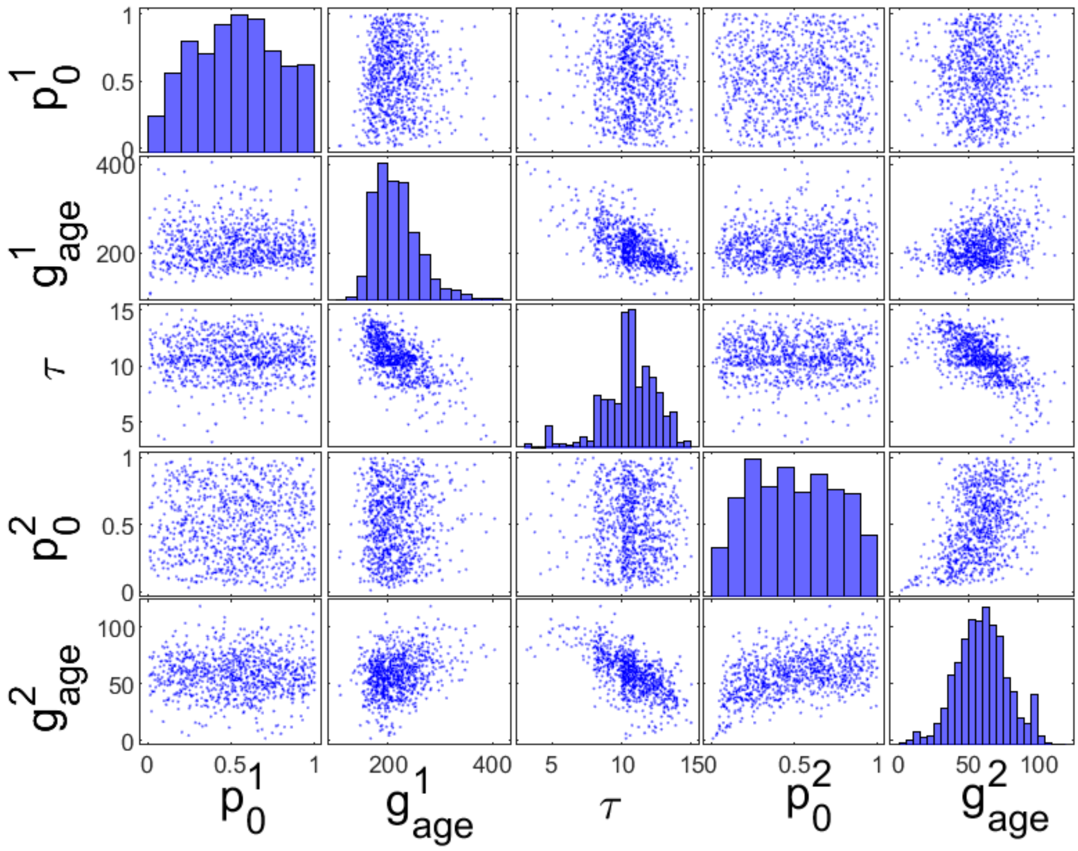

Figure S22: Bivariate plot for first mouse in pancreatic cancer dataset.

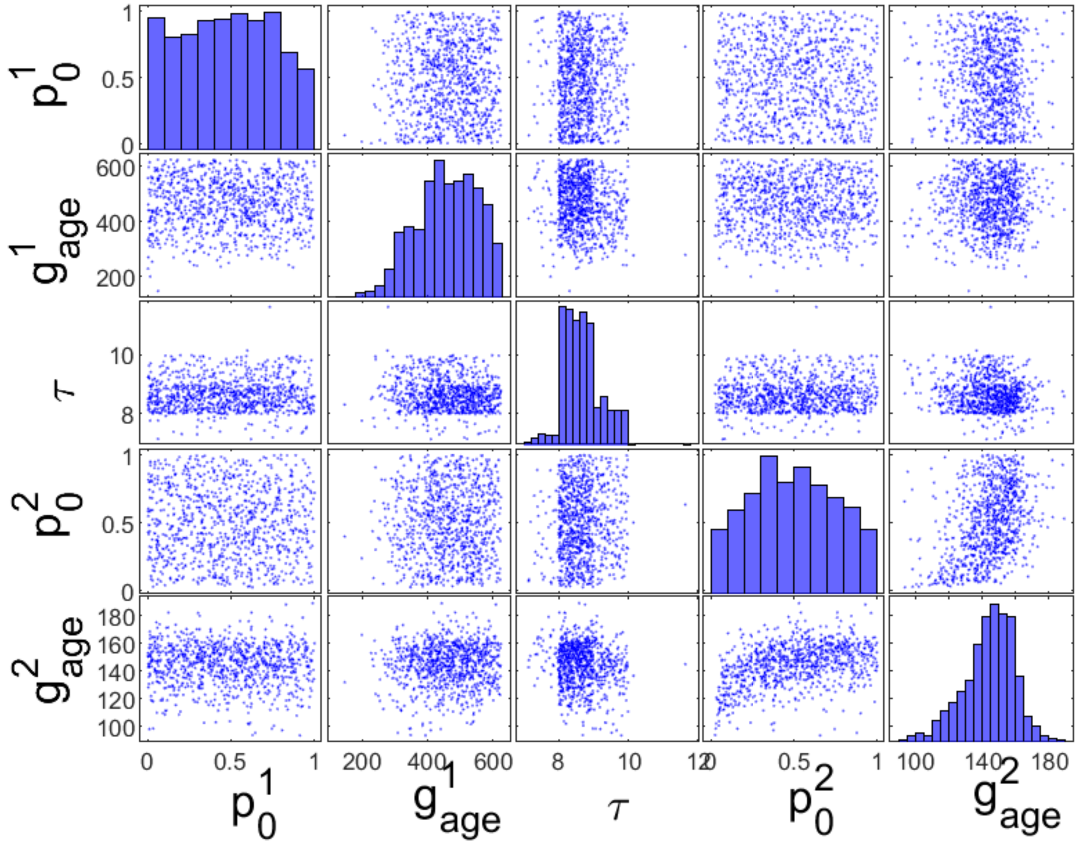

Figure S23: Bivariate plot for second mouse in pancreatic cancer dataset.

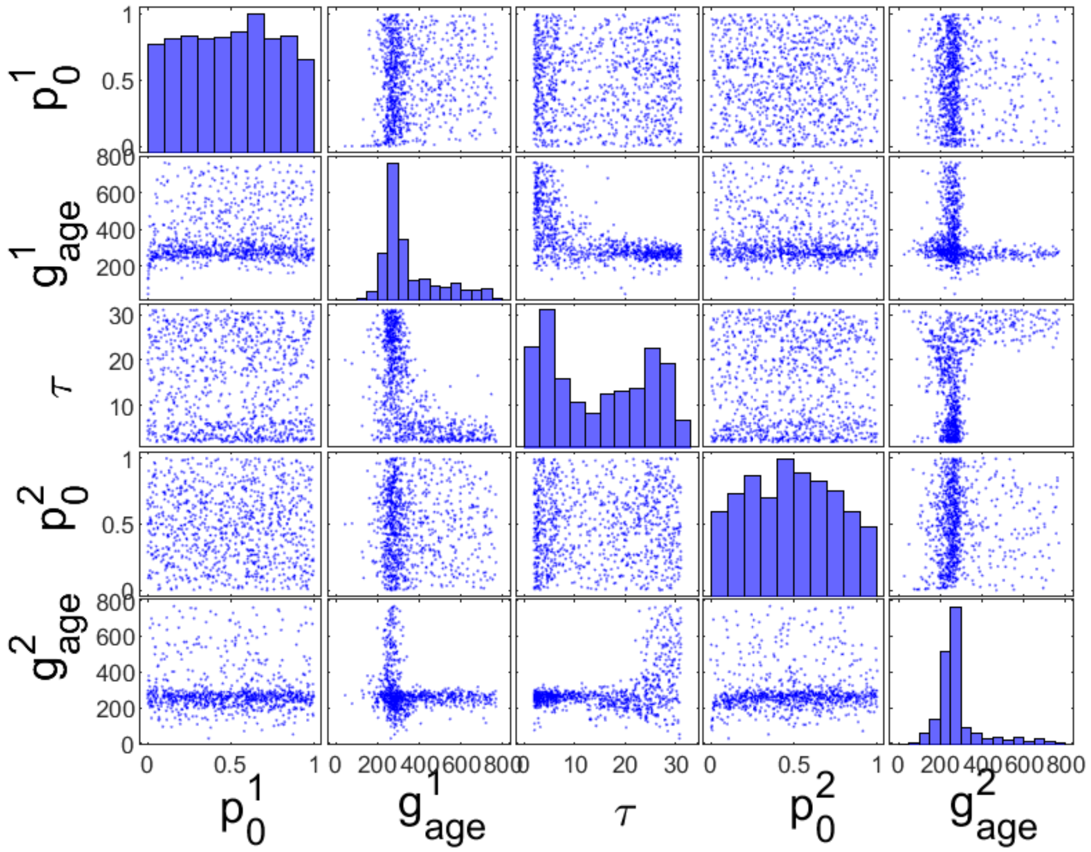

Figure S24: Bivariate plot for third mouse in pancreatic cancer dataset.

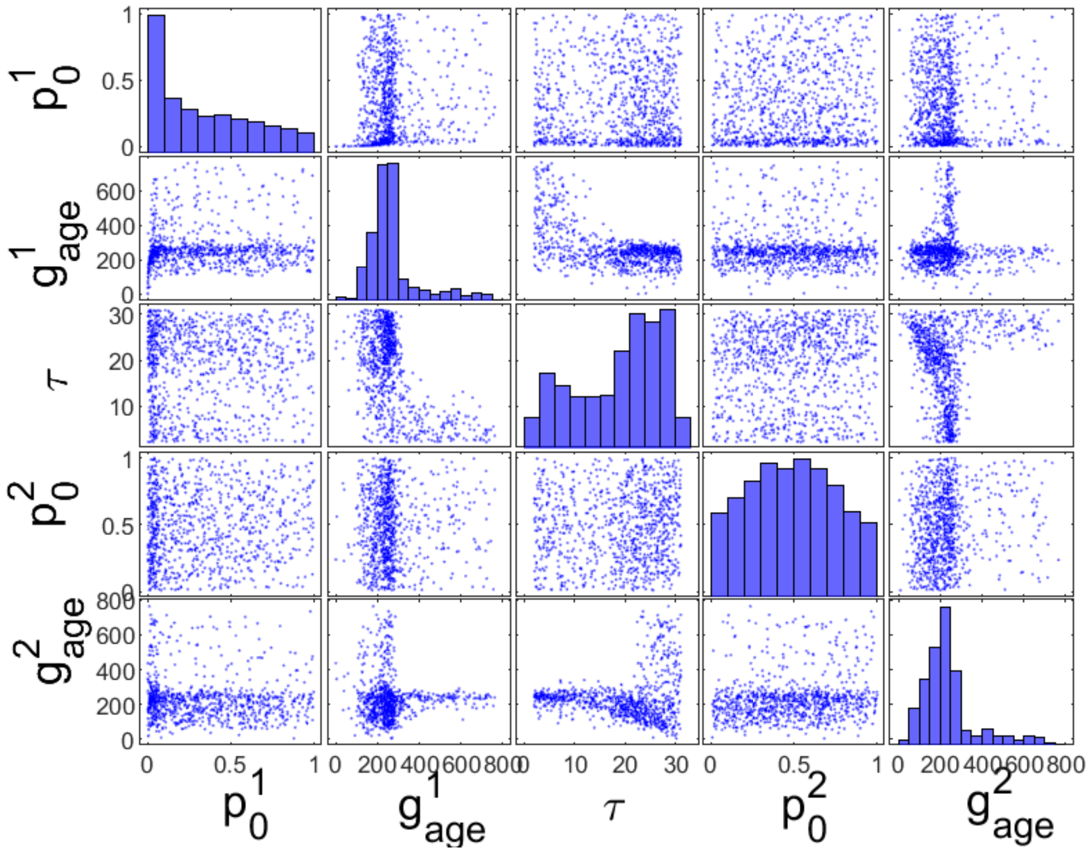

Figure S25: Bivariate plot for fourth mouse in pancreatic cancer dataset.
